# Ovarian Tumor FAK Inhibition Releases Omega-3 Fatty Acids Stimulating GATA6 Peritoneal Macrophage CXCL13 Production Enhancing Immunotherapy

**DOI:** 10.1101/2025.09.12.675975

**Authors:** Xiao Lei Chen, Kevin M Tharp, Marjaana Ojalill, Duygu Ozmadenci, Antonia Boyer, Terrance J. Hannen, Christine Lawson, Hyojae James Lee, Marvin Xia, Elise Tahon, Yichi Zhang, Cray Minor, Safir Ullah Khan, Colin C Anderson, Travis Nemkov, Michael Rose, Monica V. Estrada, Alfredo A Molinolo, Elias Warren, Patrick Penolosa, Ramez N Eskander, Michael T McHale, Shizhen E Wang, Denise C Connolly, Kathleen M Fisch, Dwayne G Stupack, David D Schlaepfer

## Abstract

High grade serous ovarian cancer (HGSOC) is the most lethal gynecologic malignancy in the USA due to chemo- and immuno-therapy resistance. We show that focal adhesion kinase (FAK) inhibition with ifebemtinib or tumor genetic FAK knockout (KO) in syngeneic ovarian tumor models stimulated resident large peritoneal macrophages to express CXCL13 chemokine and promoted B cell infiltration. Macrophage GATA6 inactivation prevented CXCL13 expression and enhanced FAK-KO tumor growth. Combining ifebemtinib with pegylated doxorubicin chemotherapy and anti-TIGIT immune checkpoint antibody extended survival with tumor-associated tertiary lymphoid structure formation. Mechanistically, FAK-KO heat-treated conditioned media contained exosomes enriched with omega-3 fatty acids which stimulated macrophage CXCL13 production. Ifebemtinib-treated tumors, FAK-KO exosomes, and purified eicosapentaenoic acid enhanced murine and human HGSOC-associated tumor macrophage reprogramming and CXCL13 expression. Overall, our studies define a tumor to macrophage signaling linkage via omega-3 exosome lipids supporting B cell recruitment, survival, immunotherapy enhancement, and actionable via small molecule FAK inhibition.

**eTOC Blurb:** High-grade serous ovarian cancer remains difficult to treat due to therapy resistance. Chen *et. al.* reveal that tumor FAK inhibition educates macrophages to express CXCL13 associated with B cell infiltration - highlighting a new therapeutic pathway linking FAK inhibition, omega-3 fatty acid containing exosomes, and macrophage mediated anti-tumor activation.

**Bullet points:**

- Genetic or small molecule FAK inhibition enhances ovarian tumor B cell infiltration
- Tumor FAK inhibition stimulates GATA6+ macrophages to make CXCL13
- FAKi, pegylated doxorubicin and anti-TIGIT promote tertiary lymphoid structures
- Omega-3 fatty acids stimulate human HGSOC ascites macrophages to make CXCL13

## INTRODUCTION

High-grade serous ovarian cancer (HGSOC) is the most common and aggressive type of ovarian cancer^1^. Surgery followed by platinum and paclitaxel chemotherapy is standard of care, but many patients develop chemotherapy resistance and succumb to disease within 5 years^2^. Platinum-resistant recurrent disease is difficult to treat and topoisomerase inhibitors such as topotecan or pegylated liposomal doxorubicin (PLD) may be used as recurrent therapies^3^. Several immune checkpoint immunotherapies (ICIs) have been tested for HGSOC, but tumor-intrinsic properties support the generation of a strong immunosuppressive tumor microenvironment (TME) which limits therapy efficacy^4^. Despite this, patients with HGSOC tumors containing elevated numbers of T cells, tumor-infiltrative B cells, and dendritic cells exhibit improved survival^5^. Thus, a greater understanding of tumor-intrinsic signals driving immunosuppression may identify new combinatorial immuno-therapeutic approaches.

Tumor-associated macrophages (TAMs) are the most abundant immune cells in HGSOC^6^. TAMs possess important regulatory immune functions via phagocytosis of cellular debris, lipid accumulation, and cytokine secretion^7^. Despite traditional macrophage roles as anti-tumoral and cytotoxic, TAMs also display considerable pro-tumor plasticity^8,9^. Analysis of TAM function in HGSOC requires consideration of distinct populations of either tissue resident large peritoneal macrophages (LPMs) expressing lineage-determining markers such as the GATA6 transcription factor or recruited small peritoneal macrophages (SPMs), derived from differentiated bone marrow-derived myeloid cells^10^. While cytotoxic chemotherapies transiently decrease total TAM populations, surviving TAMs may either support or inhibit immune surveillance^11^. Alterations in protein marker expression can indicate pro-tumor and anti-inflammatory or anti-tumor and pro-inflammatory TAM reprogramming^7^. Although GATA6^+^ LPMs are considered immune guardians of the peritoneal cavity^10,12^, the role of GATA6^+^ macrophage inactivation in murine tumor models remains unclear^13^.

In addition to important roles for TAMs and T cells in peritoneal immune surveillance, increased B cell recruitment^14,15^ and the formation of tertiary lymphoid structures (TLS) is a prognostic factor for better survival in copy-number-driven tumors like HGSOC^5,16,17^. Structurally, TLS are an ectopic collection of non-encapsulated B, T, and dendritic cells occurring at inflammation sites, including those in autoimmune disease and cancer^18^. In general, patient tumors with TLS show greater response to ICIs^19^. Mechanistically, the dialog of cytokine signaling impacting B cells and T cells in the TLS facilitates both T and B cell activation^16^. Thus, the generation of B cell recruitment signals within an HGSOC immunosuppressive TME appears critical for adaptive T cell responses^20^. Cytokines released by tumor and stromal cells also play a significant role in the formation, maturation and location of TLS within tumors^21^. In mouse ovarian cancer models, exogenous CXCL13 cytokine, also known as B cell recruitment factor, plus ICI antibodies prevented tumor growth in a CD8^+^ T cell dependent manner^22^.

The molecular mechanisms generating an HGSOC immunosuppressive TME are driven by genomic alterations^23,24^. HGSOC tumors contain mutated p53 tumor suppressor, which is observed together with several recurring regional chromosomal gains and losses^25^. Chromosomal gains at 8q24.2-8q24.3 encompass the focal adhesion (FAK) tyrosine kinase gene (*PTK2*). FAK gains are prognostic for decreased patient survival^26^ and prevalent in >75% of HGSOC tumors^27^. *PTK2* gains are associated with elevated FAK mRNA and protein levels with enhanced FAK tyrosine phosphorylation^26^. We have developed an aggressive, chemotherapy-resistant C57Bl6 syngeneic ascites-producing ovarian tumor model termed KMF (*<u>K</u>Ras*, *<u>M</u>yc*, *<u>F</u>AK* amplified) that contains DNA gains and losses in gene regions comparable to human HGSOC^26,28^. In the KMF model, genetic or small molecule FAK inhibition regulated CD155 ligand for the TIGIT (T cell immunoreceptor with Ig and ITIM domains) ICI receptor and FAK inhibition elevated TME associated CXCL13 levels in vivo^28^. However, the FAKi-stimulated cellular CXCL13 source and mediating signal(s) remain unknown.

Herein, we use ifebemtinib, a FAKi in clinical testing^29^. We find that FAKi or genetic tumor FAK inactivation increased GATA6^+^ LPM expression of CXCL13 *in vivo* associated with elevated B cell peritoneal infiltration in three independent syngeneic ovarian tumor models. FAKi combined with PLD and anti-TIGIT ICI therapy significantly increased survival with TLS formation in KMF tumor bearing mice. The FAKi CXCL13-inducing signal present in tumor conditioned medium was 56°C heat stable and associated with elevated omega-3 fatty acids levels in exosomes released from FAK-inhibited tumor cells. Notably, FAKi and the omega-3 fatty acid eicosapentaenoic acid (EPA) pushed murine LPMs toward an anti-tumor phenotype and stimulated human macrophages isolated from HGSOC patient ascites to express CXCL13. Together, we show that ovarian tumor FAK inhibition releases omega-3 fatty acids triggering GATA6 macrophage to produce CXCL13, which was associated with B cell recruitment, and the strengthening of anti-TIGIT immunotherapy.

## RESULTS

### Ifebemtinib FAK Inhibitor (FAKi) Chemotherapy Combinations in Ovarian Cancer

As pre-clinical studies have shown that FAKi combinations with paclitaxel or cisplatin can provide ovarian tumor control^26,30,31^, and platinum with paclitaxel chemotherapy is standard-of-care for HGSOC, we evaluated effects of oral ifebemtinib (FAKi) and intraperitoneal (IP) cisplatin plus (+) paclitaxel (CPT) with or without FAKi (Figure 1A) on KMF ascites-associated tumor growth and peritoneal immune cell infiltration by lavage collection followed by flow cytometry (Figure S1, and STAR Methods). Single agent FAKi reduced tumor cell number >25%, with CPT exhibiting maximal >85% tumor inhibition (Figure 1B). FAKi also increased B cell infiltration in tumor-bearing mice compared to control (P=0.042). However, whereas the combination of FAKi + CPT exhibited strong tumor control and increased T cells in the TME, this was not accompanied with elevated B cell, dendritic cell, or TAM infiltration within peritoneal ascites fluid (Figure 1B).

**Figure 1.**
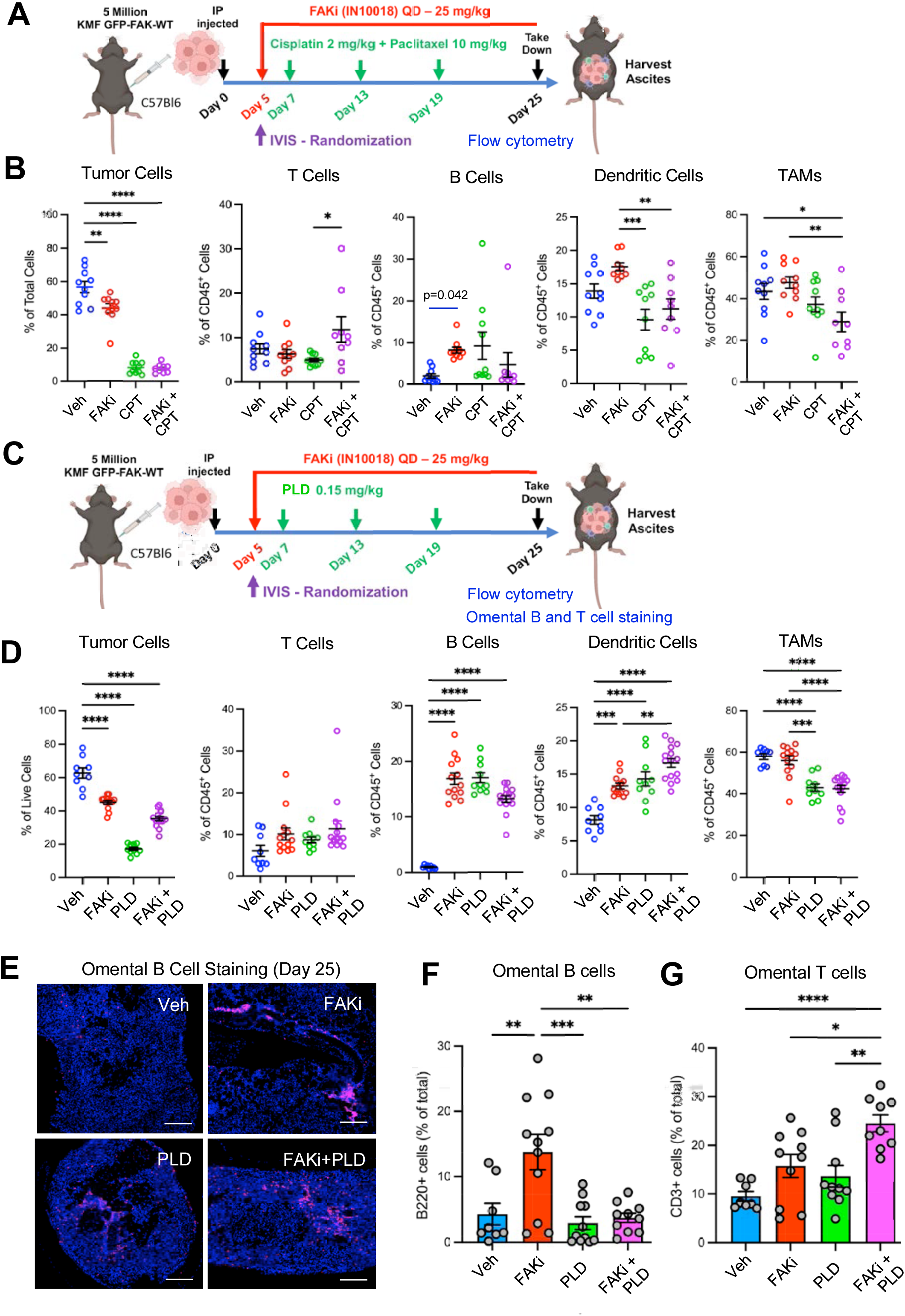
Ifebemtinib FAK Inhibitor (FAKi) with Pegylated Doxorubicin (PLD) Combine to Prevent Ovarian Tumor Growth with Enhanced B and Dendritic Cell Infiltration. (A) Experimental schematic of KMF ovarian tumor ascites model with oral FAKi and intraperitoneal (IP) cisplatin + paclitaxel (CPT) chemotherapy. (B) Flow cytometry of tumor and immune cells collected by peritoneal lavage at Day 25 and represent control (Veh, blue), FAKi (red), CPT (green), and FAKi + CPT (purple) treatments. Cell populations were defined as tumor (CD45^-^), T cell (CD45^+^ TCRβ^+^), B cell (CD45^+^ B220^+^), dendritic cell (CD45^+^ CD11c^+^), and TAM (CD45^+^ CD11b^+^ F4/80^+^). (C) Schematic of KMF tumor model treated with oral FAKi and IP PLD. (D) Flow cytometry of tumor and immune cells collected at Day 25 and represent control (blue), FAKi (red), PLD (green), and FAKi + PLD (purple) treatments. (B and D). Points from individual mice (n=9), CPT and PLD experiments were performed separately, and shown is one of two independent experiments. (E) Representative omental KMF tumor implant staining of B cells (purple) and cell nuclei (Hoechst, blue). Scale is 100 µm. (F and G) Quantitation of omental B cells (panel F) or omental T cells (panel G) from KMF tumor experiment shown in panel D. Points are from at least 8 sections from 3 mice per experimental group. (B, D, F, and G) Values are means +/- SEM (*P<0.05, **P<0.01, ***P<0.001, ****P<0.0001; one way ANOVA with Tukey’s multiple comparisons test). Panel B, T-test as noted (* P=0.042).

To test the combinatorial potential of FAKi with other chemotherapies, KMF tumor bearing mice were treated with FAKi and/or PLD (Figure 1C). As expected, FAKi reduced tumor cell number >20% with PLD exhibiting >65% tumor inhibition compared to control (Figure 1D). FAKi + PLD did not result in an additional combinatorial reduction in tumor burden. Nevertheless, FAKi, PLD, and FAKi + PLD individually and together enhanced B cell and dendritic peritoneal infiltration in tumor-bearing mice (Figure 1D). Increased B cell staining nearby omental tumor implants was also detected in FAKi-treated mice (Figures 1E and F). Although there was no significant increase in T cell number within ascites with FAKi + PLD treatment (Figure 1D), FAKi + PLD increased T cells within omental tumor implant sites (Figure 1G). Taken together, while FAKi + CPT suppressed tumor growth and increased T cell infiltration, FAKi + PLD also controlled tumor growth and promoted B and dendritic cell accumulation, indicating distinct FAKi immune-modulating effects may depend on the chemotherapeutic partner.

### Transgenic FAK KO ovarian tumor model

FAK inactivation prior to mouse adulthood can result in developmental lethality^32^. To study the effects of FAK KO in a transgenic model of ovarian cancer, FAK^fl/fl^ mice were crossed with mice expressing the SV40 T antigen (TAg^+^) under the control of the Müllerian inhibiting substance type II receptor^33^ (Figure 2A). Spontaneous ovarian tumors were isolated, cells expanded and genotyped as mouse ovarian carcinoma (MOVCAR) cells. Adenoviral (Ad) Cre recombinase was used to inactivate FAK expression *ex vivo* (Figure 2B) which did not alter PYK2 levels but resulted in E-cadherin and N-cadherin expression changes upon FAK loss (Figure 2C). There was no difference between FAK^fl/fl^ MOVCAR and Ad-Cre-treated MOVCAR FAK^-/-^ cell growth in culture (Figure 2D).

**Figure 2.**
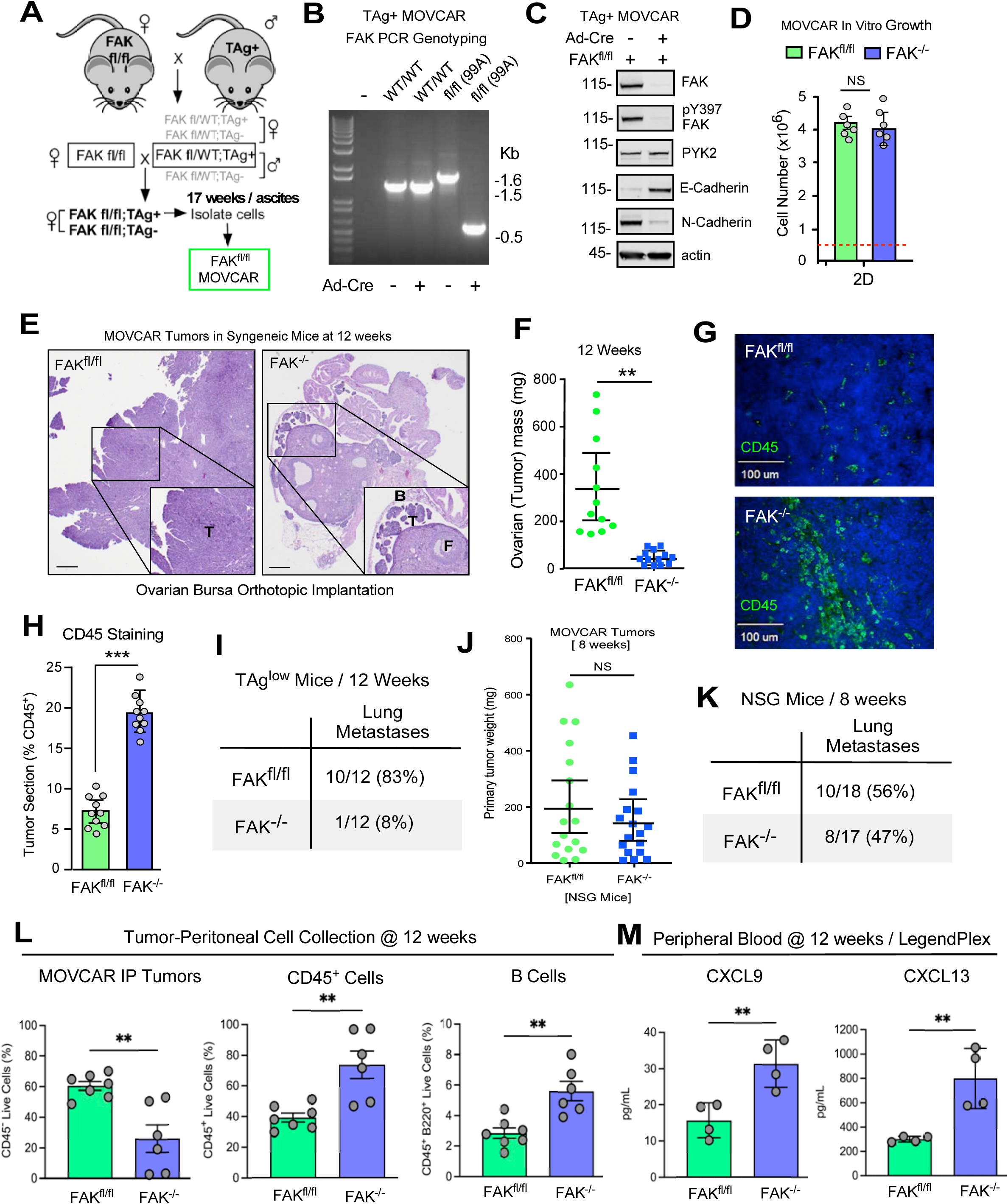
FAK KO Sensitizes Mouse Ovarian Carcinoma (MOVCAR) Tumors to TME Regulation with Elevated B Cell Infiltration and Increased Blood CXCL13. (A) Schematic summarizing FAK^fl/fl^ and SV40 T antigen (TAg) mouse crosses. Müllerian inhibiting substance type II receptor (MISIIR) promoter controls TAg expression and females form spontaneous ovarian tumors. (B) At 12 weeks, ascites cells were collected, expanded in 3D culture, and genotype determined by PCR after adenoviral Cre recombinase (Ad-Cre) transduction. Cre removal of FAK loxP DNA in MOVCAR tumor cells (FAK^-/-^) generates a 0.6 kb band. (C) Control or Ad-Cre-treated MOVCARs were immunoblotted with antibodies to FAK, pY397 FAK, PYK2, E-cadherin, N-cadherin, and actin. (D) Growth of MOVCAR FAK^fl/fl^ and FAK^-/-^ (Ad-Cre treated) cells in culture (n=6). (E) Representative H&E staining of intra-bursal implanted FAK^fl/fl^ and Ad-Cre pre-treated FAK^-/-^ MOVCAR tumors at 12 weeks. Scale is 250 µm. Inset shows magnified (2.5x) tumor (T), bursa (B), and ovarian follicle (F). (F) Mean ovarian tumor mass of implanted FAK^fl/fl^ or FAK^-/-^ MOVCARs in C57Bl6 TAg-low mice at 12 weeks. (n=12). (G) Representative staining of FAK^fl/fl^ or FAK^-/-^ MOVCAR tumors for CD45 (green) or Hoechst (blue). Scale is 100 µm. (H) Quantitation of fluorescent CD45 staining. Mean values are percent CD45-positive from 2 independent tumors (5 random fields per tumor). (I) Spontaneous lung metastasis of MOVCAR tumors from panel F. Values are percent of total mice. (J) Mean ovarian tumor mass of implanted FAK^fl/fl^ or FAK^-/-^ MOVCARs in NSG mice at 8 weeks. (n=17 or 18). (K) Spontaneous lung metastasis of MOVCAR tumors from panel J. Values are percent of total mice. (L and M) Flow cytometry of CD45^+^ and B cells of FAK^fl/fl^ or FAK^-/-^ MOVCAR ascites at 12 weeks (n=6) and cytokine quantitation of CXCL9 and CXCL13 levels in peripheral blood (n=4). (D, F, H, J, L and J) Values are means +/- SEM (**P<0.01 and ***P<0.001; T-Test). NS, not significant.

However, intrabursal injection of MOVCAR FAK^fl/fl^ cells into non-tumor prone low TAg syngeneic mice (MISIIR-TAg-Low^33^) yielded large and invasive tumors compared to MOVCAR FAK^-/-^ cells that remained as small tumors primarily within the intrabursal ovary space (Figures 2E and F). Additionally, CD45 staining of ovaries from tumor-bearing mice revealed increased immune cell presence within MOVCAR FAK^-/-^ compared to FAK^fl/fl^ tumor bearing mice (Figures 2G and H). Expectedly, decreased FAK^-/-^ MOVCAR tumor burden was associated with few spontaneous lung metastases (Figure 2I). Conversely, in immune-deficient mice, there was no suppression of FAK^-/-^ MOVCAR growth or progression to lung metastasis (Figures 2J and K), underscoring the importance of TME suppressive effects on MOVCAR FAK^-/-^ cells in immunocompetent mice. Analysis of IP cell fractions revealed decreased tumor cells but significantly increased CD45^+^ immune cells in syngeneic MISIIR-TAg-Low mice engrafted with MOVCAR FAK^-/-^ cells compared to MOVCAR FAK^fl/fl^ cells (Figure 2L). Decreased FAK^-/-^ MOVCAR tumor burden was associated with increased B cell infiltration and elevated CXCL9 and CXCL13 cytokine levels in peripheral blood (Figures 2L and M). Together, our results support the notion that tumor FAK loss or inhibition may elicit signals that impact immune TME and that FAK^-/-^ MOVCARs are sensitive to repressive TME signals.

### Ifebemtinib and PLD Increase Large Peritoneal Macrophage (LPM) CXCL13 Expression

Our previously published studies showed that VS4718 FAKi increased *Cxcl13* mRNA in non-tumor cells of the KMF TME^28^. To determine the FAKi-stimulated cellular source of CXCL13, flow cytometry and single cell RNA sequencing (scRNAseq) was performed with ifebemtinib FAKi, PLD, or FAKi + PLD in KMF tumor bearing mice (Figure 3). In FAKi-treated mice, stimulated CXCL13 expression was highest in CD45^+^ CD11b^+^ F4/80^high^ LPMs compared to CD45^+^ CD11b^+^ F4/80^low^ SPMs or CD45^+^ CD11b^-^ F4/80^-^ myeloid cells (Figures 3B and C). Interestingly, PLD treatment also increased CD45^+^ CD11b^+^ F4/80^high^ LPM CXCL13 expression but no combinatorial effect with FAKi + PLD on CXCL13 levels was observed (Figure 3C) which was consistent with the lack of FAKi + PLD effects to boost B cell infiltration further (Figure 1D). scRNAseq analyses confirmed that FAKi, PLD, and FAKi + PLD increased TME B cells compared to vehicle control as represented by Uniform Manifold Approximation and Projection (UMAP) dimensional reduction plots (Figure 3D). *Cxcl13* mRNA expression was elevated in myeloid cells by FAKi, PLD, and FAKi + PLD treatment of tumor-bearing mice (Figures 3E and Table S1). Strong induction of *Cxcl13* mRNA expression was detected in myeloid but not B or T/NK cell populations (Figure 3F) by FAKi, PLD, and FAKi + PLD. UMAP sub-clustering of myeloid cells revealed population-level changes in mRNA expression upon FAKi, PLD, or PLD + FAKi treatment of KMF tumor-bearing mice (Figure 3G). Notably, FAKi-induced *Cxcl13* expression partially overlapped with *Gata6* and *Tim4* mRNAs (Figure 3H), which are markers of LPMs^34^. Moreover, TCGA analyses revealed that HGSOC tumor mRNAs associated with CXCL13 include CXCL9 chemokine, TIGIT, and other T cell markers (Table S2). Taken together, our results support the notion that FAKi and PLD increase CXCL13 in LPMs of the TME with *Gata6*⁺ or *Timd*4⁺ LPMs as the putative primary cellular source.

**Figure 3.**
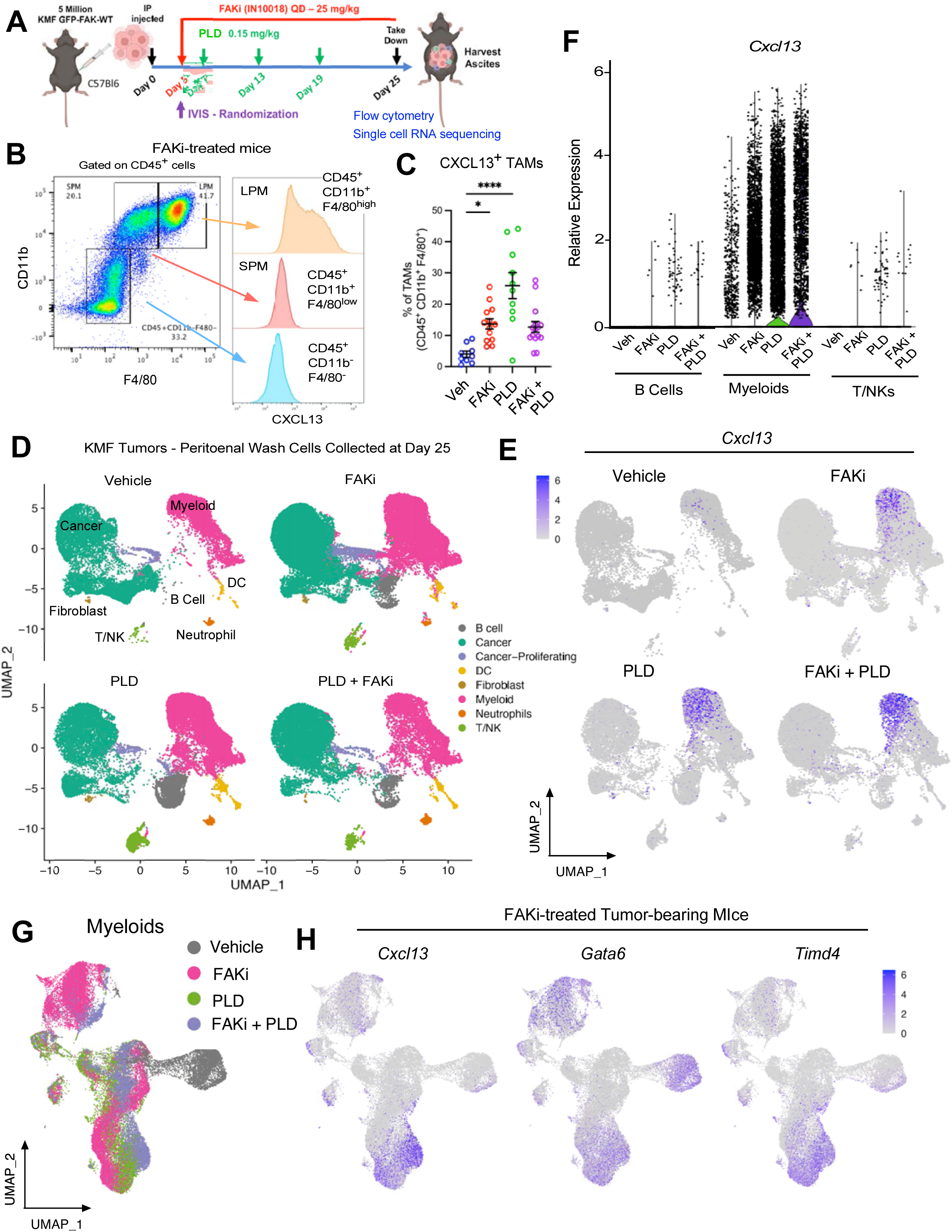
Flow Cytometry and Single Cell Sequencing of KMF Peritoneal TME Reveal FAKi- and PLD-induced Large Peritoneal Macrophage Associated CXCL13 Expression. (A) Schematic of KMF ovarian tumor model treated with oral FAKi and IP PLD. (B) Representative flow cytometry of FAKi-treated KMF tumor-bearing mice of CD45^+^ CD11b^+^ F4/80^high^ LPM TAMs, CD45^+^ CD11b^+^ F4/80^low^ small peritoneal macrophages (SPM), and CD45^+^ CD11b^-^ F4/80^-^ immune cells. (C) Flow cytometry of CD45^+^ CD11b^+^ F4/80^+^ CXCL13^+^ TAMs from vehicle, FAKi, PLD, and PLD + FAKi treated mice bearing KMF tumors isolated at Day 25. Values are means +/- SEM (*P<0.05 and ***P<0.001; one way ANOVA with Tukey’s multiple comparisons test; n=10). (D) UMAP dimensionality reduction plot of single cell RNA sequencing (scRNAseq) performed on peritoneal tumor and immune cells collected Day 25. Shown are composite cell identifications denoted by color (legend at right) for all experimental groups. (E) UMAP plot (single color) denoting CXCL13 mRNA expressing cells in vehicle, FAKi, PLD, and FAK + PLD experimental groups at Day 25. Scale at left. (F) Relative CXCL13 mRNA expression in B cells, myeloid, or T/NK cells in KMF tumor bearing mice as determined by scRNAseq and separated by experimental groups. (G) UMAP plot of CD45^+^ myeloid cells as determined by scRNAseq showing changes in cell subpopulations in vehicle, FAKi, PLD, and FAK + PLD experimental groups at Day 25. (H) UMAP plot (single color) of myeloid cells expressing *Cxcl13*, Gata6, and *Timd4* mRNA in FAKi-treated experimental group at Day 25. Scale at right.

### FAKi with PLD Strengthen anti-TIGIT Immunotherapy and Promote Survival

As FAKi and PLD enhance B cell infiltration in the TME, and FAKi reduced CD155 ligand for TIGIT on KMF cells^28^, we tested whether anti-TIGIT inhibitory antibody as part of a triple chemo- and immuno-therapy combination over 24 days may maximally enhance B and T cell TME recruitment (Figure 4A). Anti-TIGIT alone had negligible effects on KMF tumor burden or changes in immune cell infiltration compared to IgG control (Figure 4B). TIGIT + PLD potently reduced tumor cells and resulted in a TME that was enriched in B cells but not CD8^+^ T cells (Figure 4B). TIGIT + FAKi reduced tumor burden at Day 25, dramatically reduced CD155 TIGIT ligand on KMF cells, and increased B cells in the TME compared to control. Interestingly TIGIT + FAKi increased dendritic cell TME infiltration compared to anti-TIGIT alone (Figure 4B). TIGIT + FAKi + PLD resulted in tumor control, tumor CD155 reduction, elevated B cells, increased CD8^+^ T cells, and greater dendritic cell TME recruitment compared to single or double therapies (Figure 4B).

**Figure 4.**
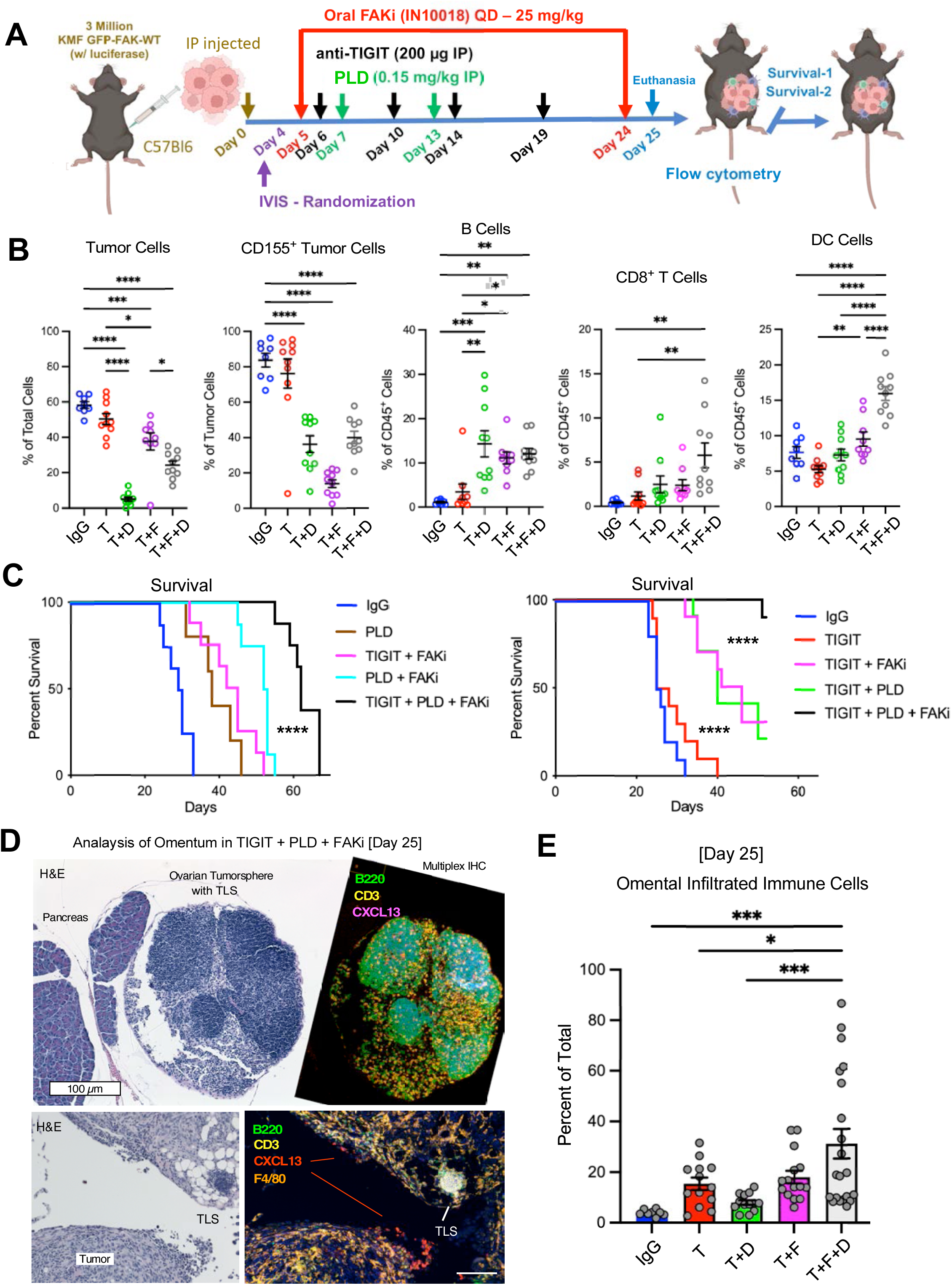
Combination FAKi and PLD Chemotherapy Strengthen anti-TIGIT Immunotherapy and Promote Extended Survival with TLS Formation. (A) Schematic of KMF tumor model testing single agent or combinations of FAKi, PLD, and anti-TIGIT. Treatments were stopped at Day 24, and in separate experiments, cells were harvested-analyzed at Day 25 (panel B), or mice were evaluated for survival (panel C). (B) Flow cytometry of vehicle (V, blue), TIGIT (T, red), TIGIT + PLD (T+D, green), TIGIT + FAKi (T+F, purple), and TIGIT + PLD + FAKi (T+D+F, clear) experimental groups (n=9). Cell populations were defined as tumor (CD45^-^ and CD155^+^), B cells (CD45^+^ B220^+^), CD8 T cells (CD45^+^ TCRβ^+^ CD8^+^) and dendritic cells (CD45^+^ CD11c^+^). Points are individual mice (n=9). (C) KMF tumor-bearing mouse survival. Experimental groups (n=10) match panel B with addition of PLD-only and PLD + FAKi. (B and C) Data shown are from one of two independent experiments. (D) Serial tissue section from omental tumor implants stained by H&E and multiplex IHC with antibodies to B cells (B220, green), T cells (CD3, yellow), and CXCL13 (magenta). Scale bar is 100 µm. (E) Quantitation of infiltrated immune cell clusters within omentum at Day 25. Points are from non-overlapping tumor sections and immune cells values presented as percent of total area. (B, C, and E) Values are means +/- SEM (*P<0.05, **P<0.01, ***P<0.001, ****P<0.0001; one way ANOVA with Tukey’s multiple comparisons test).

In independent experiments, KMF tumor bearing mice were treated with the same 24-day protocol, treatment stopped, and extended survival measured (Figures 4A and C). All control mice reached a humane endpoint by Day 35. Anti-TIGIT alone did not extend survival. PLD-only mice exhibited tumor regrowth that limited survival to 40-45 days, and PLD + FAKi treated mice exhibited survival to 50 days (Figure 4C). Significantly greater survival (>60 days) occurred in TIGIT + PLD + FAKi treated mice (Figure 4C). In a second independent experiment, dual combinations TIGIT + FAKi and TIGIT + PLD yielded equivalent survival beyond 50 days (Figure 4C). However, TIGIT + PLD + FAKi treatment extended survival significantly greater than either dual drug combinations (Figure 4C). As FAKi + PLD is being tested in a clinical trial for recurrent HGSOC [NCT06014528], our results support the future combinatorial testing of ICI immunotherapies such as anti-TIGIT with FAKi + PLD chemotherapies.

### Tertiary Lymphoid Structure (TLS) Formation with FAKi, PLD, and anti-TIGIT Treatments

The molecular mechanisms supporting TLS formation remain under investigation^19^. As immune cell populations collected and analyzed in the peritoneal TME can differ from solid tumors, T and B cell staining of omental KMF tumor implants was performed (Figure S2A). Control tumors had few immune cells, PLD alone increased T cell infiltration, FAKi treatment resulted in B cell enriched sites within tumors, and PLD + FAKi promoted tumor-associated T and B cell small clusters. Increased T and B cell infiltration of solid tumors with PLD + FAKi treatment correlated with increased survival (Figure 4C).

Although anti-TIGIT alone did not impact survival or T cell tumor infiltration in the KMF model, TIGIT + PLD and TIGIT + FAKi significantly extended survival (Figure 4C) and resulted in elevated immune cell infiltration of tumors as measured at Day 25 (Figure S2B). Importantly, mouse survival beyond 60 days was associated with omental immune cell infiltration and TLS formation upon TIGIT + PLD + FAKi therapy (Figures 4D, E and Figure S2B). Additionally, omental tumor staining detected F4/80^+^ CXCL13^+^ TAMs nearby tumor-associated clusters of B and T cells in tumor-bearing TIGIT + PLD + FAKi treated mice (Figure 4D). Taken together, our results show that FAKi and PLD chemotherapy strengthen anti-TIGIT immunotherapy with enhanced B cell recruitment, survival and TLS formation.

### Tumor FAK Expression and Activity Control Macrophage CXCL13 Expression and B cell TME Recruitment

To define better the role of FAK protein and/or activity in tumor cells, we employed clonal KMF FAK-KO cells generated by CRISPR and made comparisons to KMF FAK-KO cells stably reconstituted with FAK-WT or a kinase-inactive FAK-K454R point mutant. FAK-WT and FAK-R454 proteins were equally expressed (Figure S3A), with FAK-WT cells producing significantly greater tumor burden than FAK-KO and FAK-R454 cells within 25 days (Figures S3B and C). scRNAseq was performed on tumor and immune cells obtained by peritoneal lavage at Day 25 and a notable UMAP difference between groups was the presence of B cells in the FAK-KO and FAK-R454 TME compared the FAK-WT TME (Figure 5A). Interestingly, this pattern of B cell distribution paralleled myeloid-associated CXCL13 expression in the FAK-KO and FAK-R454 but not the FAK-WT TME (Figure 5B). To extend these results, we used CRISPR to inactivate FAK or the related PYK2 kinase expression in murine HGS2 fallopian tube derived tumor cells from p53^-/-^ PTEN^-/-^ and BRCA2^-/-^ mice (Figure S3D). Loss of FAK but not PYK2 inhibited HGS2 syngeneic tumor growth (Figure S3E) and flow cytometry analyses showed that FAK but not PYK2 loss inhibited HGS2 CD155 TIGIT ligand expression *in vivo* (Figure S3F). Analysis of CD45^+^ F4/80^+^ CD11b^+^ TAMs showed significantly increased CXCL13^+^ TAMs from FAK^-/-^ compared to PYK2^-/-^ and parental HGS2 tumor-bearing mice (Figure S3G). Taken together, three independent ovarian syngeneic tumor models link tumor FAK inactivation with decreased tumor burden, increased TAM CXCL13 production and enhanced B cell TME infiltration.

**Figure 5.**
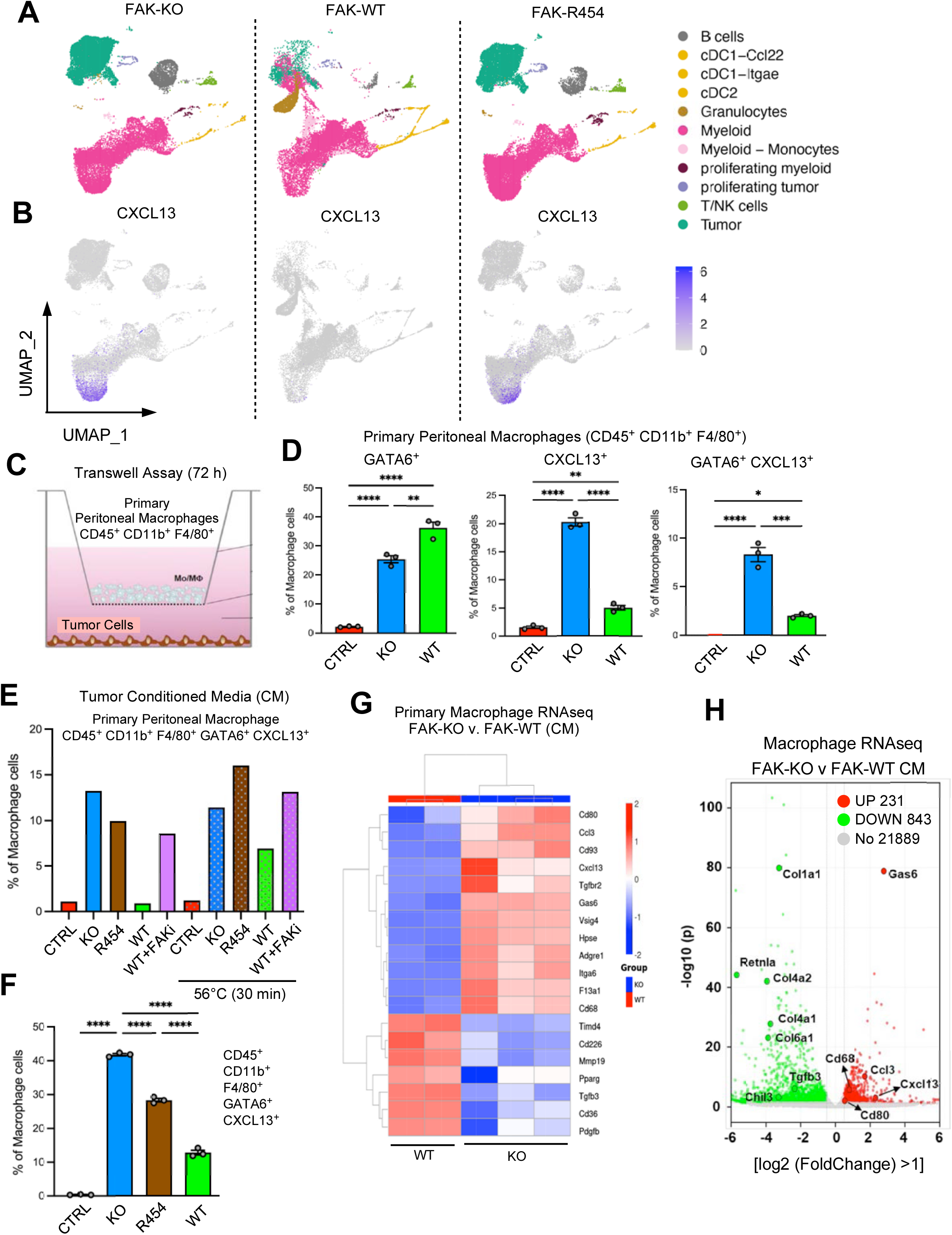
Tumor FAK Inactivation Releases a Heat-stable Macrophage Polarization and CXCL13-inducing Factor into Conditioned Media (CM) (A) UMAP plot of scRNAseq performed on peritoneal cells from KMF FAK-KO, FAK-WT, and FAK R454 tumor bearing mice at Day 25. Shown are composite cell identifications denoted by color (legend at right). (B) UMAP plot (single color) denoting *Cxcl13* mRNA expressing cells in FAK-KO, FAK-WT, and FAK R454 tumor bearing mice at Day 25. Scale at right. (C) Schematic of transwell KMF tumor cell (lower chamber) with primary C57Bl6 peritoneal macrophage (upper chamber) co-culture signaling assay. (D) Flow cytometry quantitation of peritoneal CD45^+^ CD11b^+^ F4/80^+^ macrophages for changes in GATA6 and CXCl13 expression in response to control (CTRL), FAK-KO, or KMF FAK-WT CM. (E) Shown is percentage of CD45^+^ CD11b^+^ F4/80^+^ GATA6^+^ CXCL13^+^ macrophages after incubation with control (CTRL) or FAK-KO, FAK-WT, FAK-R454, and FAK-WT cells pre-treated with FAKi (1 µM) CM. In parallel, CM was heated (56°C, 30 min) and macrophage responses measured by flow cytometry. (F) Quantitation of heat-treated FAK-KO, FAK-R454, and FAK-WT CM stimulated peritoneal macrophage CXCL13 expression. D and F) Values are means +/- SEM from three independent co-culture samples per experimental group (*P<0.05, **P<0.01, ***P<0.001, ****P<0.0001 by ANOVA with Tukey post hoc test). (G) RNAseq heatmap of elevated (red) or decreased (blue) peritoneal macrophage mRNAs after incubation with FAK-KO or FAK-WT CM. (H) Volcano plot of differentially expressed macrophage mRNAs upon incubation with FAK-KO or FAK-WT CM.

### Tumor FAK Inhibition Releases a Heat-stable CXCL13-inducing and Macrophage Anti-Tumor Reprogramming Factor

To determine if FAK-KO induced signals promoting TAM CXCL13 expression *in vivo* were mediated by cell-cell contact or via an indirect soluble mediator, KMF FAK-KO or FAK-WT cells were incubated with unstimulated C57Bl6 myeloid-macrophages isolated by peritoneal lavage (Figure 5C). FAK-KO but not FAK-WT co-culture stimulated macrophage (CD45^+^ CD11b^+^ F4/80^+^) CXCL13 expression (data not shown), and this was reproduced in cell-separated transwells (Figure 5D). Interestingly, both FAK-KO and FAK-WT cells in transwells stimulated increased macrophage GATA6^+^ expression but only FAK-KO cells boosted CXCL13 and increased double-positive GATA6^+^ CXCL13^+^ macrophages compared to signals generated by FAK-WT cells (Figure 5D). The mediator of macrophage CXCL13 induction was a soluble factor present in serum-free conditioned media (CM) of FAK-KO and FAK-R454, but not FAK-WT cells (Figure 5E). Heat treatment at 56°C to denature tumor-released proteins and cytokines in serum-free conditions did not alter FAK-KO or FAK-R454 CM-enhanced macrophage CXCL13 expression (Figure 5F). This heat-stable factor was also generated by pre-treating FAK-WT cells with FAKi prior CM collection, indicating that FAK inhibition was "stimulating" the release of this reprogramming factor (Figure 5E).

This tumor to macrophage signaling linkage was conserved as murine PMJ2-R macrophages also responded with increased CXCL13 expression when exposed to heat-treated FAK-KO KMF or FAK^-/-^ HGS2 CM (Figure S4). Notably, FAKi-treated KMF-WT or FAKi-treated HGS2 cells also generated CM that could stimulate PMJ2-R CXCL13 expression. To determine tumor CM-induced macrophage mRNA changes, C57Bl6 peritoneal macrophages were incubated with heat-treated KMF FAK-KO or FAK-WT CM and analyzed by bulk RNA sequencing Figure 5G (Table S3). Differential expression analyses revealed elevated levels of *Cxcl13, Adgre1, Fgfr1, Itga6, Vsig4, Itgb1, and Lrg1* mRNAs which were part of the KMF tumor-bearing *Cxcl13* myeloid population identified by scRNAseq (Table S1, grouping 2, highlighted). In total, 843 macrophage mRNAs were downregulated and 231 were upregulated (log_2_ fold change > 1) when comparing FAK-WT versus FAK-KO CM effects (Figures 5G and H). Anti-tumor macrophage markers (*Gas6*, *CD80, CD68, Ccl3*) were upregulated in the presence of FAK-KO CM, while pro-tumor macrophage markers (*Retnla, Chil3, Tgfb3, Col1a1, Col6a1, and Col12a1*) were elevated in FAK-WT CM treated macrophages and verified by Q-RT-PCR (Figure S5). In total, our results support the notion that soluble heat-stable signals released from FAK-KO tumor cells increase CXCL13 and push peritoneal macrophages toward an anti-tumor phenotype.

### Macrophage GATA6 Knockout Prevents FAK-KO Induced CXCL13 Expression and Augments FAK-KO Tumor Growth

As scRNAseq showed that FAKi-associated macrophage *Cxcl13* mRNA expression overlapped with *Gata6* in myeloid-specific analyses (Figure 3H), we used LysM Cre macrophage-specific inactivation of GATA6^fl/fl^ mice to test the impact of GATA6 loss on KMF FAK-KO CM-stimulated CXCL13 expression (Figure 6A). LysM Cre^+^ GATA6^fl/fl^ mice exhibit a selective loss of resident LPMs but not macrophages in other organs^34^. Quantitative PCR confirmed GATA6 mRNA reduction from total peritoneal lavage-isolated macrophages from LysM Cre^+^ GATA6^fl/fl^ (GATA6-KO) compared to LysM Cre^-^ GATA6^fl/fl^ (GATA6-WT) mice (Figure 6B). FAK-KO but not FAK-WT CM stimulated CXCL13 expression from GATA6-WT but not GATA6-KO (CD45^+^ CD11b^+^ F4/80^+^) macrophages (Figures 6C and D).

**Figure 6.**
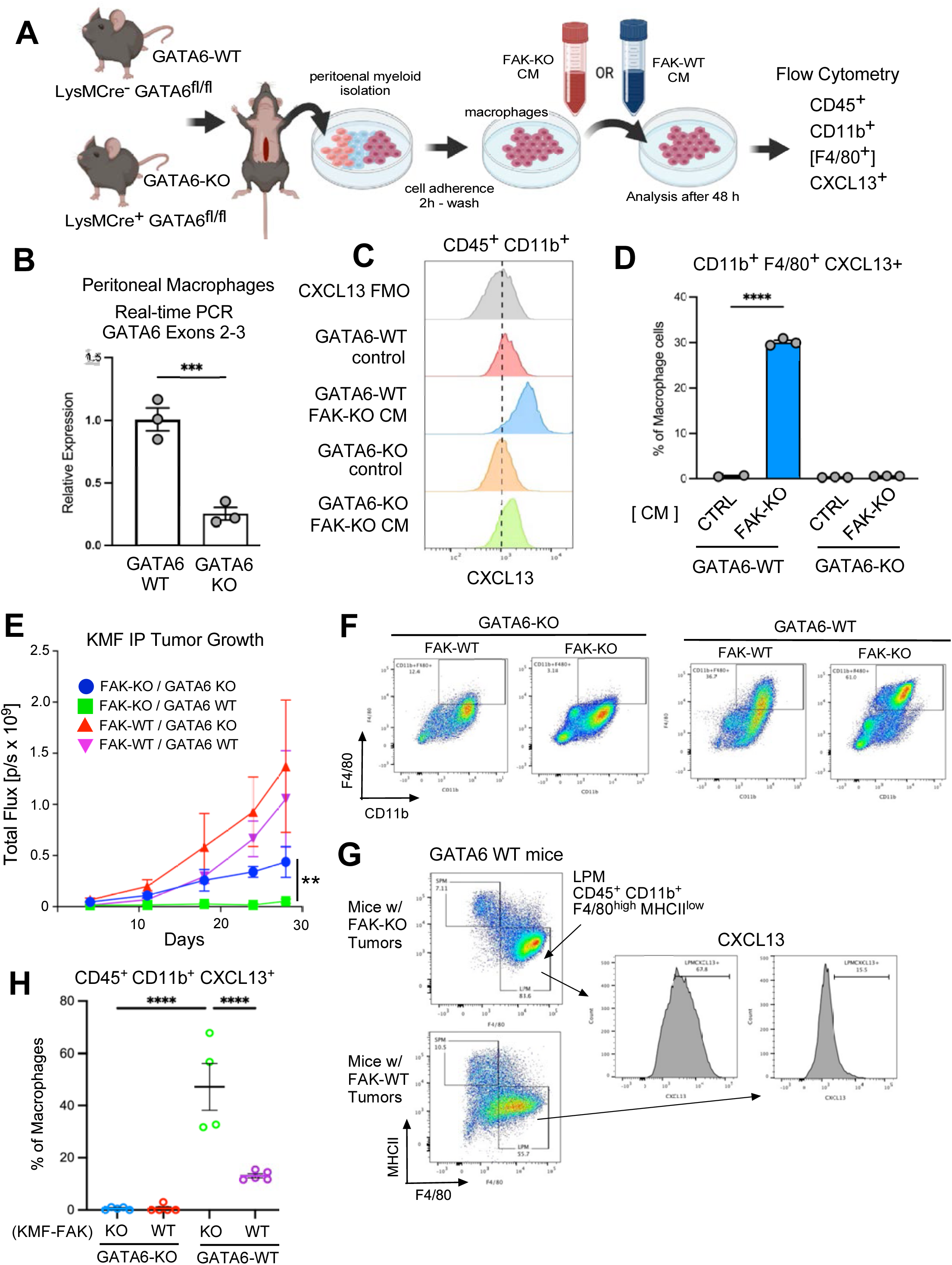
Macrophage GATA6 Knockout Prevents FAK-KO Induced CXCL13 Expression with Augmented FAK-KO Syngeneic Tumor Growth. (A) Schematic of peritoneal macrophage isolation from LysMCre^-^ GATA6^fl/fl^ YFP^+^ (GATA6-WT) and LysMCre^+^ GATA6^fl/fl^ YFP^+^ (GATA6-KO) mice and stimulation with heat-treated FAK-KO or FAK-WT CM. (B) Quantitative RT-PCR of *Gata6* exons 2-3 from total peritoneal macrophages isolated from unstimulated GATA6-WT and GATA6-KO mice. Values are means +/- SEM (***P<0.001 by unpaired T test). (C) Representative flow cytometry of induced CXCL13 expression in CD45^+^ CD11b^+^ F4/80^+^ peritoneal GATA6-WT but not GATA6-KO macrophages by FAK-KO CM. (D) Quantitation of FAK-KO CM stimulated CXCL13 expression in GATA6-WT macrophages *in vitro*. (E) 3 million KMF FAK-KO or FAK-WT were IP-injected in GATA6-KO and GATA6-WT mice and monitored for tumor growth by IVIS (n=5 per group). (F) Representative flow cytometry CD45^+^ CD11b^+^ F4/80^+^ macrophage profiles from tumor-bearing GATA6-KO and GATA6-WT mice. (G) Representative CD45^+^ CD11b^+^ F4/80^high^ MHC-II^low^ flow cytometry profiles showing levels of FAK-KO or FAK-WT tumor associated CXCL13 expression. (H) Quantitation of CD45^+^ CD11b^+^ CXCL13 expression. (D, E and I) Values are means +/- SEM (**P<0.01, ****P<0.0001 by ANOVA with Tukey post hoc test).

To determine the effect of GATA6 loss on syngeneic tumor growth, KMF FAK-KO and FAK-WT tumor growth were compared upon injection into GATA6-WT and GATA6-KO mice (Figure 6E). As expected, KMF FAK-WT cells generated greater tumor burden than FAK-KO cells in both GATA6-WT and GATA6-KO mice. Surprisingly, KMF FAK-KO cells exhibited enhanced tumor growth in GATA6-KO compared to GATA6-WT mice (Figure 6E). Additionally, peritoneal TAMs from GATA6-KO mice exhibited low F4/80 surface expression (also observed in unstimulated GATA6-KO mice, data not shown) compared to CD45^+^ CD11b^+^ F4/80^+^ TAMs from GATA6-WT mice (Figure 6F). CXCL13 was expressed by CD45^+^ CD11b^+^ F4/80^high^ MHCII^low^ LPMs present in GATA6-WT mice (Figure 6G) and a significantly greater percent of GATA6-WT LPMs expressed CXCL13 in tumor-bearing FAK-KO compared to FAK-WT mice (Figure 6H). Together, these results support the notion that LysMCre^+^ GATA6 inactivation decreased LPMs, prevented FAK-KO tumor-induced CXCL13 expression, and was associated with augmented FAK-KO tumor growth.

### CXCL13 Induction by Omega-3 Fatty Acid Enriched Exosomes from FAK-KO CM

Building on the finding that the CXCL13 inducing factor in CM was heat stable, we analyzed intact cells (Figure 7A) and methanol-extracted CM from KMF FAK-KO and FAK-WT cells by lipid mass spectrometry (Figure 7B, and Tables S4 and S5). Several differences were detected, and α-linoleic acid (αLA), an omega−6 fatty acid and eicosapentaenoic acid (EPA), an omega−3 fatty acid were elevated in FAK-KO compared to FAK-WT CM (Figure 7B). EPA is a dietary acquired omega-3 fatty acid, is heat stable, found in salmon and fish oil supplements, and known for its potential heart health benefits^35^. To determine if these lipids could impact C57Bl6 LPMs *in vitro*, purified αLA or EPA were added to FAK-WT CM and assayed for CXCL13-promoting activity (Figure 7C). EPA but not αLA addition potently induced CXCL13 expression in CD45^+^ CD11b^+^ F4/80^+^ LPMs. Using human THP-1 macrophages, EPA and the omega-3 fatty acid docosahexaenoic acid (DHA) significantly increased *Cxcl13* mRNA levels within 18 h (Figure 7D). These results suggest omega-3 fatty acids may be signaling lipids in FAK-KO CM.

**Figure 7.**
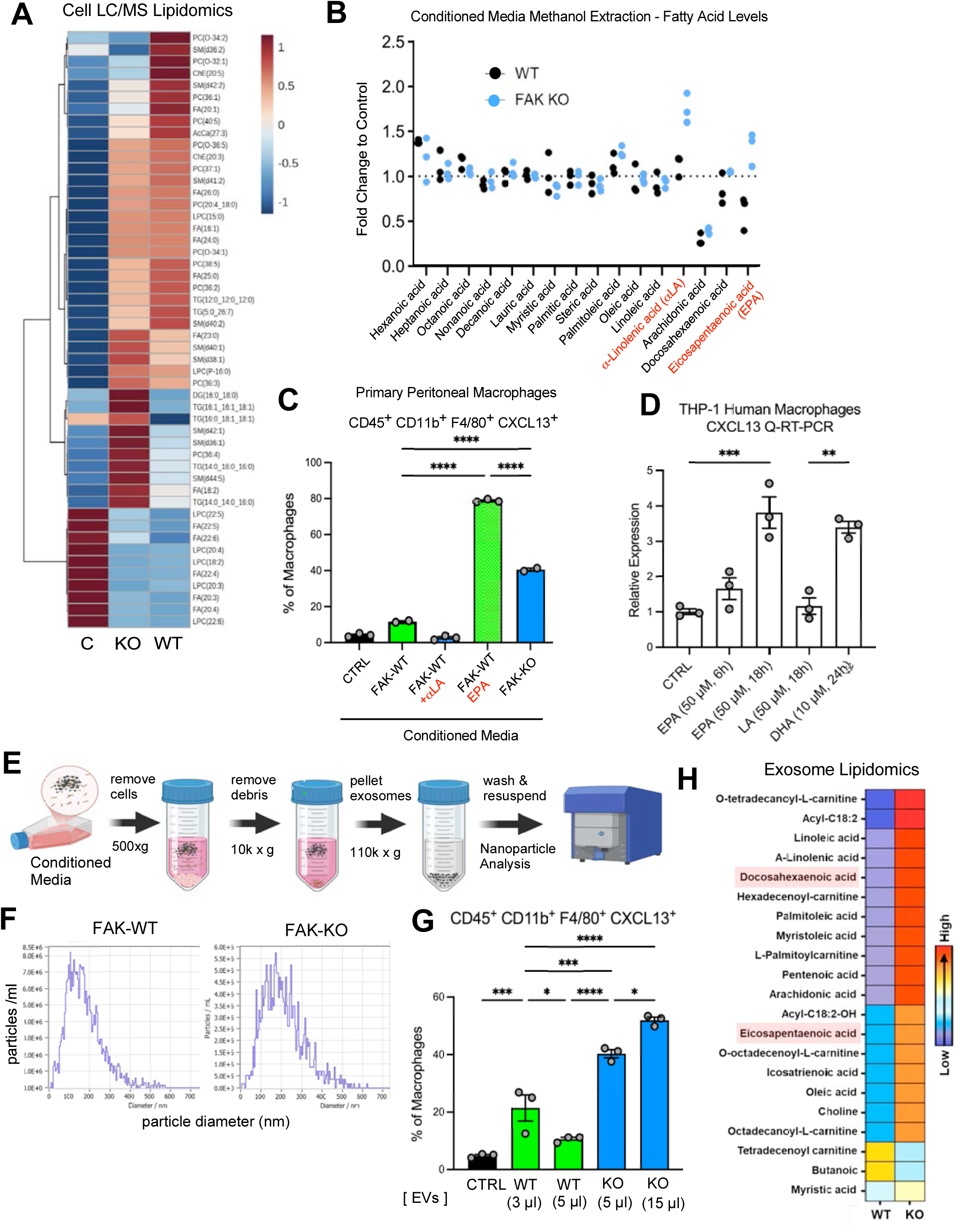
Exosome-associated Omega-3 Fatty Acids from FAK-KO Tumor Cells Stimulate Macrophage CXCL13 Expression. (A) Heatmap of cellular lipid differences in FAK-KO and FAK-WT KMF cells. C, control is serum-free cell media. (B) Mass spectrometry quantification of CM lipids from FAK-KO (blue circles) and FAK-WT (black circles) cells. Values (n=3) are fold-change compared to control. α-linolenic acid (αLA) and eicosapentaenoic acid (EPA) in FAK-KO media are highlighted in red. (C) Quantitation of CXCL13 in CD45^+^ CD11b^+^ F4/80^+^ peritoneal macrophages upon stimulation with FAK-WT CM, FAK-WT CM with 0.01% purified αLA, with added EPA, and compared to FAK-KO CM. D) EPA and docosahexaenoic acid (DHA) but not α-linolenic (αLA) stimulate CXCL13 mRNA expression in THP-1 human macrophages. (C and D). Values are means +/- SEM (**P<0.01, ***P<0.001, ****P<0.0001; one way ANOVA with Tukey’s multiple comparisons test, n=3). (E) Schematic summary of exosome purification from serum-free FAK-KO and FAK-WT CM by repeated ultracentrifugation. (F) Nanoparticle tracking profile of purified FAK-KO and FAK-WT exosomes. (G) Percentage of CD45^+^ CD11b^+^ F4/80^+^ CXCL13^+^ murine peritoneal macrophages by flow cytometry after control (PBS) or increasing volumes of purified FAK-KO or FAK-WT exosomes. (C, D, and G). Values are means +/- SEM (**P<0.01, ***P<0.001, ****P<0.0001; one way ANOVA with Tukey’s multiple comparisons test, n=3). (H) Heatmap differences of mass spectrometry lipidomic analyses of purified FAK-KO or FAK-WT exosomes. DHA and EPA are highlighted.

In aqueous solutions, lipids can form structures like micelles and bilayers. Exosomes are secreted lipid bilayer-coated vesicles ranging in size from 30 to 150 nm that play important roles in cell-to-cell signaling within the TME^36^. Exosomes from FAK-KO and FAK-WT CM were purified by repeated centrifugation steps, suspended in PBS, and analyzed by nanoparticle tracking (Figure 7E). Starting from the same number of cells, purified FAK-WT and FAK-KO exosomes were approximately the same final concentration with particle size ranging from 50 to 300 nm (Figure 7F). Notably, FAK-KO but not FAK-WT exosomes induced CD45^+^ CD11b^+^ F4/80^+^ LPMs to make CXCL13 (Figure 7G) and contained elevated levels of DHA and EPA omega-3 fatty acids (Figure 7H and Table S6). Together, these results show that tumor FAK inactivation releases exosomes with elevated level of omega-3 fatty acids that stimulate LPMs to make CXCL13.

### CXCL13 Induction by FAKi or EPA Using Cells from Human Ovarian Tumor Ascites

Abdominal paracentesis is used to relieve ovarian cancer patients of malignant ascites, which is a mix of tumor and immune cells that can be evaluated *ex vivo* (Figure 8A and Table S7). Human macrophages are identified by specific surface markers (CD14 and CD68) and macrophages can be polarized toward anti-tumor (elevated CD86) or pro-tumor (elevated CD163) phenotypes (Figure 8B). Incubation of total ascites cells with FAKi or EPA significantly increased (CD45^+^ CD14^+^ CD68^+^) CD86^+^ TAM markers and decreased (CD45^+^ CD14^+^ CD68^+^) CD163^+^ surface TAM markers (Figures 8C and D). This increase in (CD45^+^ CD14^+^ CD68^+^) CD86^+^ TAMs was paralleled by increased CXCL13 expression (Figure 8E). Using gradient-purified TAMs from patient ascites, EPA increased *Cxcl13* mRNA (Figure 8F). EPA-stimulated CXCL13 was produced by CD45^+^ CD14^+^ CD86^+^ but not CD45^+^ CD14^+^ CD86^-^ TAMs (Figure 8G) with EPA and DHA omega-3 fatty acids showing equivalent CXCL13-inducing activities (Figure 8H). Taken together, our results show that FAKi effects on tumors and EPA effects on TAMs promote an anti-tumor murine and human peritoneal macrophage phenotype with increased CXCL13 expression.

**Figure 8.**
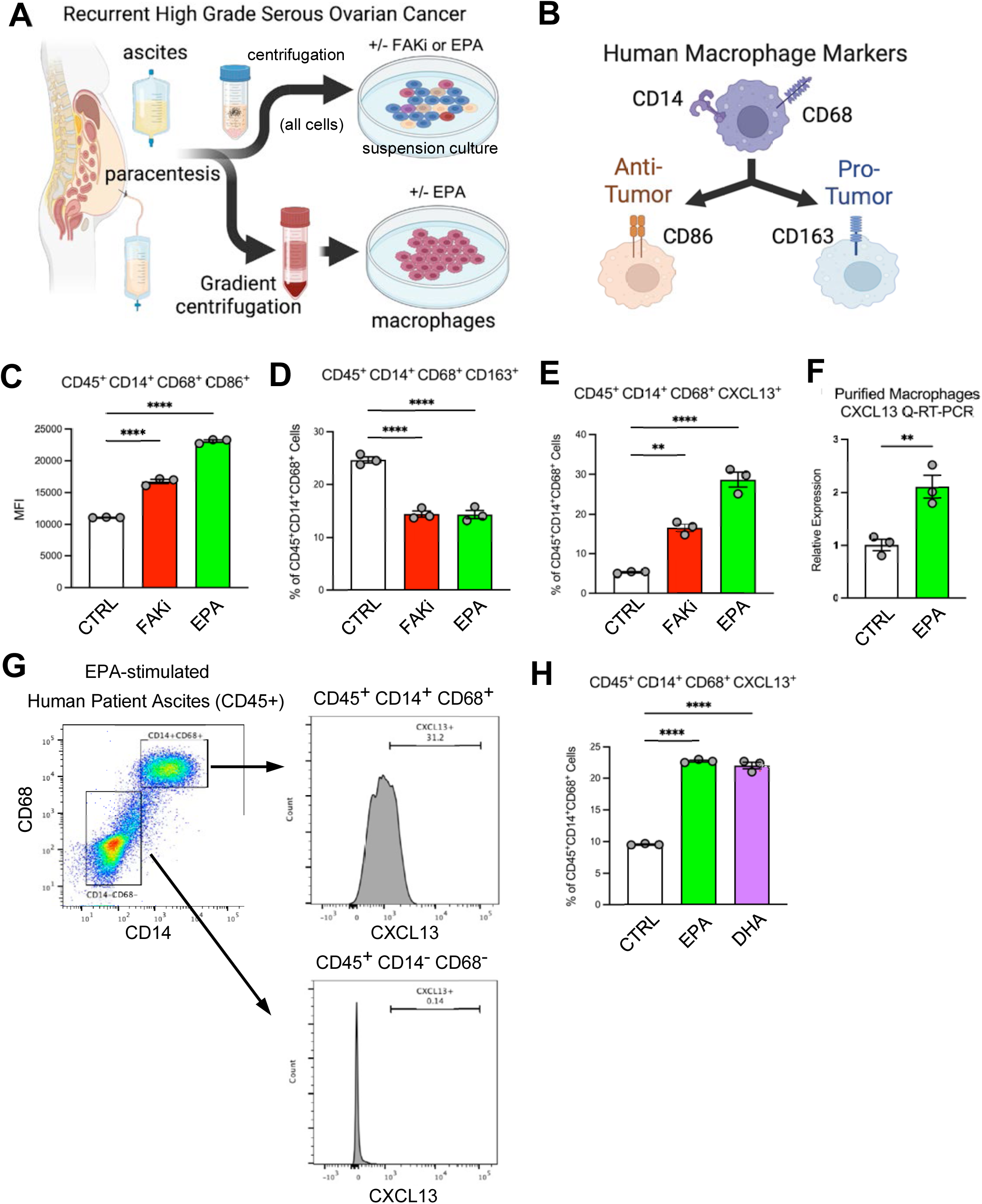
Human Ascites Macrophages Produce CXCL13 After FAKi-Tumor or Omega-3 Fatty Acid Stimulation. (A) Schematic of ovarian can patient ascites collection by paracentesis, total cell or gradient-purified macrophage isolation, and evaluation of effects of FAKi or EPA addition after 48 h. (B) Simplified model of surface protein markers that define human macrophages (CD14^+^ and CD68^+^) and either anti-tumor (CD86^+^) or pro-tumor (CD163) polarization. (C) Flow cytometry quantitation of CD45^+^ CD14^+^ CD68^+^ CD86^+^ macrophages upon FAKi (1 µM) or purified EPA (50 µM) addition. (D) Quantitation of CD45^+^ CD14^+^ CD68^+^ CD163^+^ macrophages as in panel C. (E) Quantitation of CD45^+^ CD14^+^ CD68^+^ CXCL13^+^ macrophages as in panel C. (F) Quantitative real-time reverse-transcriptase PCR of CXCL13 mRNA levels in EPA-stimulated purified patient macrophages. (G) Representative flow cytometry gating of EPA-stimulated CD45^+^ CD14^+^ CD68^+^ macrophage CXCL13 expression. (H) Percentage of CD45^+^ CD14^+^ CD68^+^ CXCL13-expressing macrophages upon EPA or DHA stimulation. (C, D, E, F and H). Values are means +/- SEM (**P<0.01, ***P<0.001, ****P<0.0001; one way ANOVA with Tukey’s multiple comparisons test, n=3). CTRL represents no addition to base media.

## DISCUSSION

The small molecule FAKi defactinib in combination with the RAF-MEK inhibitor avutometinib has received accelerated FDA approval for recurrent low grade serous ovarian cancer^37^. However, HGSOC is a molecularly different tumor with p53 mutations and FAK copy number gains without RAS-MAPK pathway mutational activation. Moreover, pre-clinical tumor models show that small molecule FAK inhibition can impact both tumor and stromal cells of the TME^38,39^. Increased complexity associated with interpretations of FAKi action are due in part to different tumor types and the use of pre-clinical FAKi compounds that inhibit both FAK and the related PYK2 kinase.

Herein, we used ifebemtinib, a FAK but limited PYK2 inhibitor, that is being tested in combination with PLD in a Phase II trial for recurrent platinum-resistant HGSOC [NCT06014528]. Using multiple syngeneic ovarian tumor models, we revealed that tumor intrinsic FAK loss, genetic FAK kinase inactivation or oral ifebemtinib FAKi treatment promoted B cell infiltration into the peritoneal TME associated with the release of omega-3 fatty acid enriched exosomes from FAK-inhibited tumor cells. FAKi-triggered release of exosomes is a lipid-associated signal to LPMs to make CXCL13, also known as B cell recruitment factor. Moreover, increased levels of CXCL13 could be detected in the peripheral blood of mice with FAK^-/-^ compared FAK^+/+^ tumors. Overall, our studies with mice and human tumor ascites revealed that ovarian tumor FAK inhibition educates macrophages to express CXCL13 - highlighting a new therapeutic pathway linking FAK inhibition, omega-3 fatty acid containing exosomes, and macrophage reprogramming toward an anti-tumor phenotype.

Surprisingly, we found that FAKi immune-modulating effects are impacted by the chemotherapeutic partner. Whereas the combination of FAKi with cisplatin and paclitaxel provided strong tumor inhibition, beneficial effects on the TME were limited. However, combinations of FAKi and PLD exhibited tumor control as well as facilitating recruitment of B cells and dendritic cells into the peritoneal tumor TME. Interestingly, the addition of an anti-TIGIT ICI antibody did not inhibit tumor growth or alter ovarian TME by itself, but adding anti-TIGIT to FAKi + PLD chemotherapy triggered maximal ovarian tumor inhibition, elevated B cells, increased CD8^+^ T cells, greater dendritic cell infiltration, TLS formation, and significantly longer survival.

CXCL13 plays a critical role in orchestrating T and B cell interactions essential for maturation of the immune response^40,41^. CXCL13 is recognized as a positive prognostic indicator in HGSOC^22^, and in this context, it was expected that the CXCL13 source would be follicular T cells^42^. Surprisingly, flow cytometry and single cell sequencing show that GATA6^+^ tissue resident LPMs are the FAKi-stimulated CXCL13 source in mice. This differs from other studies noting that ovarian tumor cells can produce CXCL13 upon CDK4/6 inhibition^43^ and that FAK-inhibition alters cytokine expression from other tumors^44^. In fact, macrophage-targeted knockout of GATA6 was associated with reduced CXCL13 levels and the increased growth of FAK^-/-^ tumors, showing that TME effects are limiting to FAK-null tumor growth *in vivo*.

Tumor cells can uptake omega-3 fatty acids from the environment^45^. This is especially true for HGSOC that typically spreads to the omentum, an adipocyte-rich tissue that contributes to circulating fatty acid levels^46^. Omega-3 fatty acids like docosahexaenoic acid (DHA) and eicosapentaenoic acid (EPA) are best known for supporting cardiovascular health, but EPA also possesses anti-tumor activity and is in clinical trials for colon cancer^47^. Our studies add the growing literature on the positive role of omega-3 fatty acids in supporting an immune-favorable TME^48^. Overall, our mechanistic studies define a new tumor to macrophage signaling relationship via exosome lipids influencing B cell infiltration of the TME that is therapeutically actionable via small molecule FAK inhibition.

### Limitations of the Study

Our analyses with HGSOC patient tumor samples collected by paracentesis was limited in sample number but revealed a potential tumor to macrophage signaling response that may be independent of prior chemotherapy treatments or tumor genetic-limiting response. In addition, as GATA6 expression in macrophages differs from that in mice^49^, our parallel findings with human ovarian TAMs ascites supports the notion that tumor-FAKi and EPA-macrophage signaling is a conserved linkage. Our results also complement earlier studies showing that GATA6 macrophages regulate gut IgA production through peritoneal B1 cells^50^ and that CXCL13 is required for B1 cell homing, antibody production, and body cavity immunity^51^. Accordingly, several unknowns exist in this signaling linkage, start with the molecular mechanism(s) of FAKi initiated release of omega-3 enriched exosomes as well as the potential lipid receptor(s) on macrophages and the signal which increase CXCL13 mRNA. Future studies using our mouse ovarian syngeneic models will provide foundational insights into how FAK inhibition impacts tumor bioactive lipid exosome content, signaling to GATA6 macrophages, B cell infiltration and maturation, with signals that normalize the TME and strengthen immunotherapy.

## Supporting information

Key Resource Table

## Author Contributions

Conceived and designed the study: DGS and DDS

Designed the experiments: XL, KT, DGS, and DDS

Provided technical and experimental support: MO, AB, TJH, CL, MX, ET, JZ, CM, and SEW

Performed the scRNA-seq experiments: DO, XL

Performed bioinformatic analyses: KMF, HJL, and SUK

Performed IHC: MR

Performed pathological assessment: MVE, AAM

Provided mouse models and expertise: DCC

Collected human samples and related clinical data: EW and PP

Provided clinical expertise: RNE and MTM

Discussed and interpreted the data: XL, KT, MO, CL, KMF, DGS, and DDS

Wrote the manuscript with all authors’ contribution: DDS

Edited the manuscript: DGS, KMF, DCC, and DDS

## RESOURCE AVAILABILITY

### Lead contact

Further information and requests for materials should be directed to the lead contact, David Schlaepfer.

### Materials availability

All materials used in this study are available from the lead contact with a completed materials transfer agreement.

### Data and code availability

The total and single cell RNA sequencing FASTQ files have been deposited to the National Library of Medicine Sequence Read Archive under the accession numbers SRRXXXXXX.

## STAR METHODS

Detailed methods are provided in the online version of this paper and include the following:

## ACKNOLWEDGMENTS

We thank InxMed Inc. for Ifebemtinib, Nine Girls Ask? for the purchase of equipment used in this study, and Gwendalyn J. Randolph (Washing University School of Medicine) for the LysMcre GATA6^fl/fl^ YFP^+^ mice. We appreciate insights provided by Dr. Hui Chen and Dr. Judith A. Varner at UCSD on murine macrophage biology.

## Titles of Supplemental Figures

**Figure S1.**
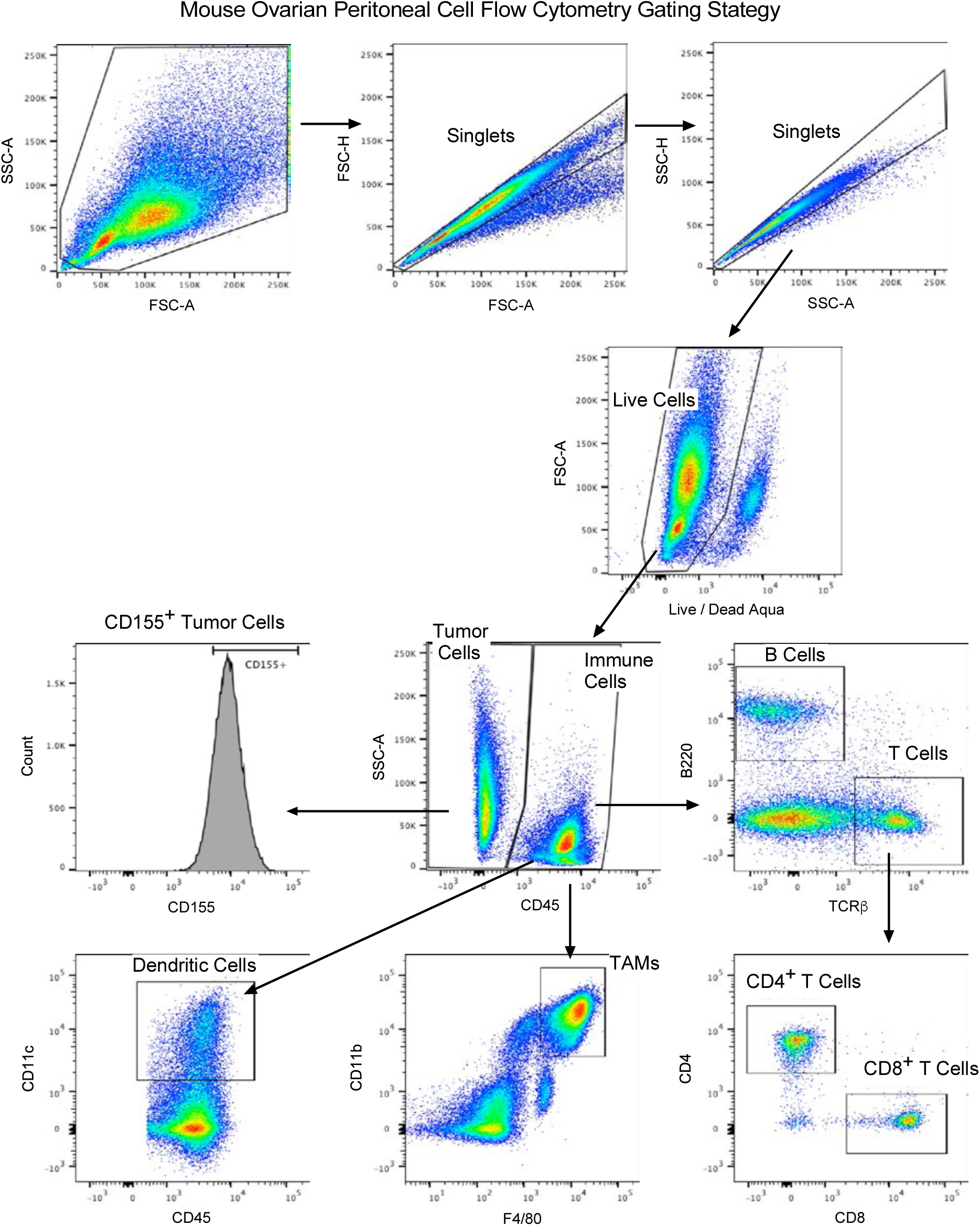
Murine Tumor Ascites Flow Cytometry Gating Strategy. Related to STAR Methods Tumor (CD45^-^) were separated from CD45^+^ immune cells, and sub-populations of peritoneal immune cells were analyzed using validated antibodies. Peritoneal wash from tumor-free mice was used to establish immune cell gating parameters.

**Figure S2.**
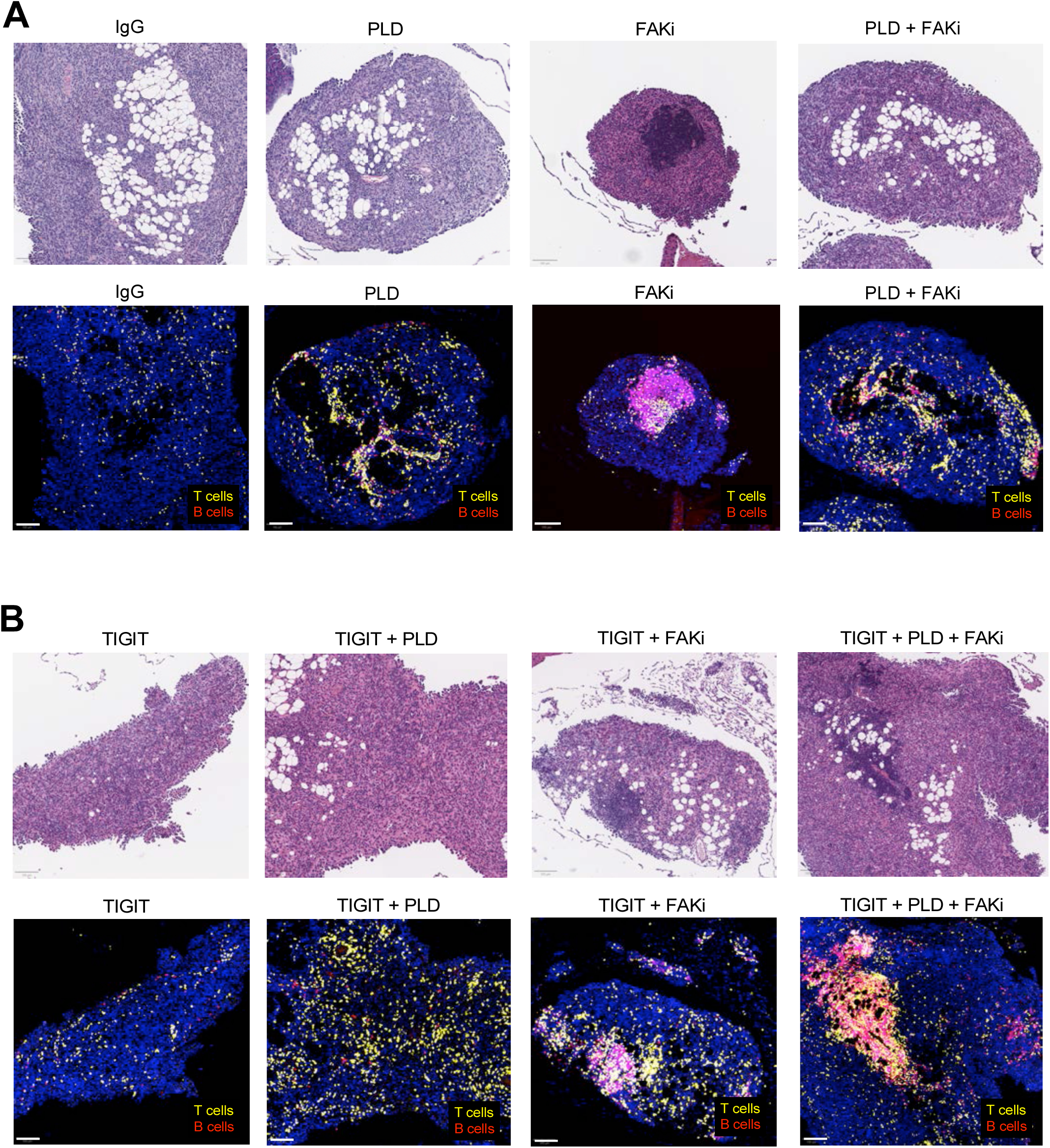
Infiltrated Immune Cells in Omentum of KMF Tumor-bearing Mice. Related to Figure 4 (A and B) Sequential FFPE omental tumor sections were analyzed by H&E and multiplex for T cells (yellow) and B cells (red) plus nuclei (blue). Scale is 100 µm. (A) Representative staining from the indicated experimental groups at Day 25. Related to Figure 1D. (B) Representative staining from the indicated experimental groups at Day 25. Related to Figure 4B. KMF Syngeneic Ovarian Tumor Model

**Figure S3.**
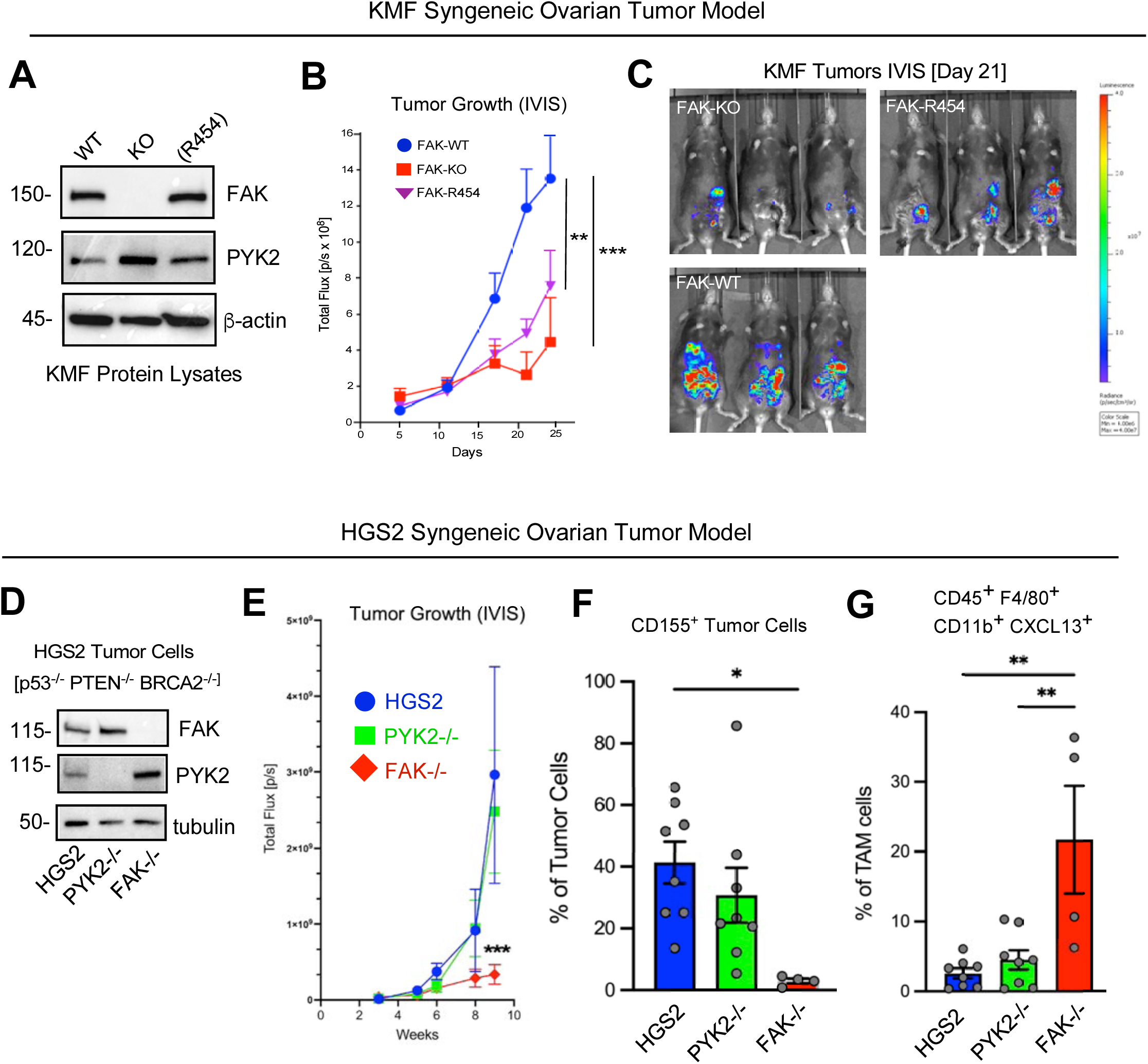
FAK Knockout in KMF and HGS2 Syngeneic Tumor Models. Related to Figure 5. (A) Immunoblotting of KT13 FAK KO KMF cells reconstituted with wildtype (WT) or kinase-dead (R454) FAK as GFP fusion proteins. FAK-related PYK2 and actin blotting as controls. (B) 5 million of the indicated KMF cells were evaluated by IP tumor growth by IVIS imaging. (C) Representative IVIS images of FAK-KO, FAK-R454, and FAK-WT KMF tumor bearing mice at Day 21. Scale bar at right. (D) Immunoblotting of parental, PYK2, and FAK knockout HGS2 ovarian tumor cells. Tubulin is loading control. (E) 5 million of the indicated HGS2 cells were evaluated by IP tumor growth by IVIS imaging. (F) Acistes tumor cell enumeration and flow cytometry of CD45^-^ CD155^+^ HGS2 tumor cells for the indicated experimental groups. (G) Flow cytometry quantitation of CD45^+^ F4/80^+^ Cd11b^+^ CXCL13^+^ macrophages in HGS2 tumor-bearing mice. (B, E-G) Values are means +/- SEM (*P<0.05, **P<0.01, ***P<0.001; one way ANOVA with Tukey’s multiple comparisons test, n=8. (F and G) HGS2 FAK^-/-^ n=4.

**Figure S4.**
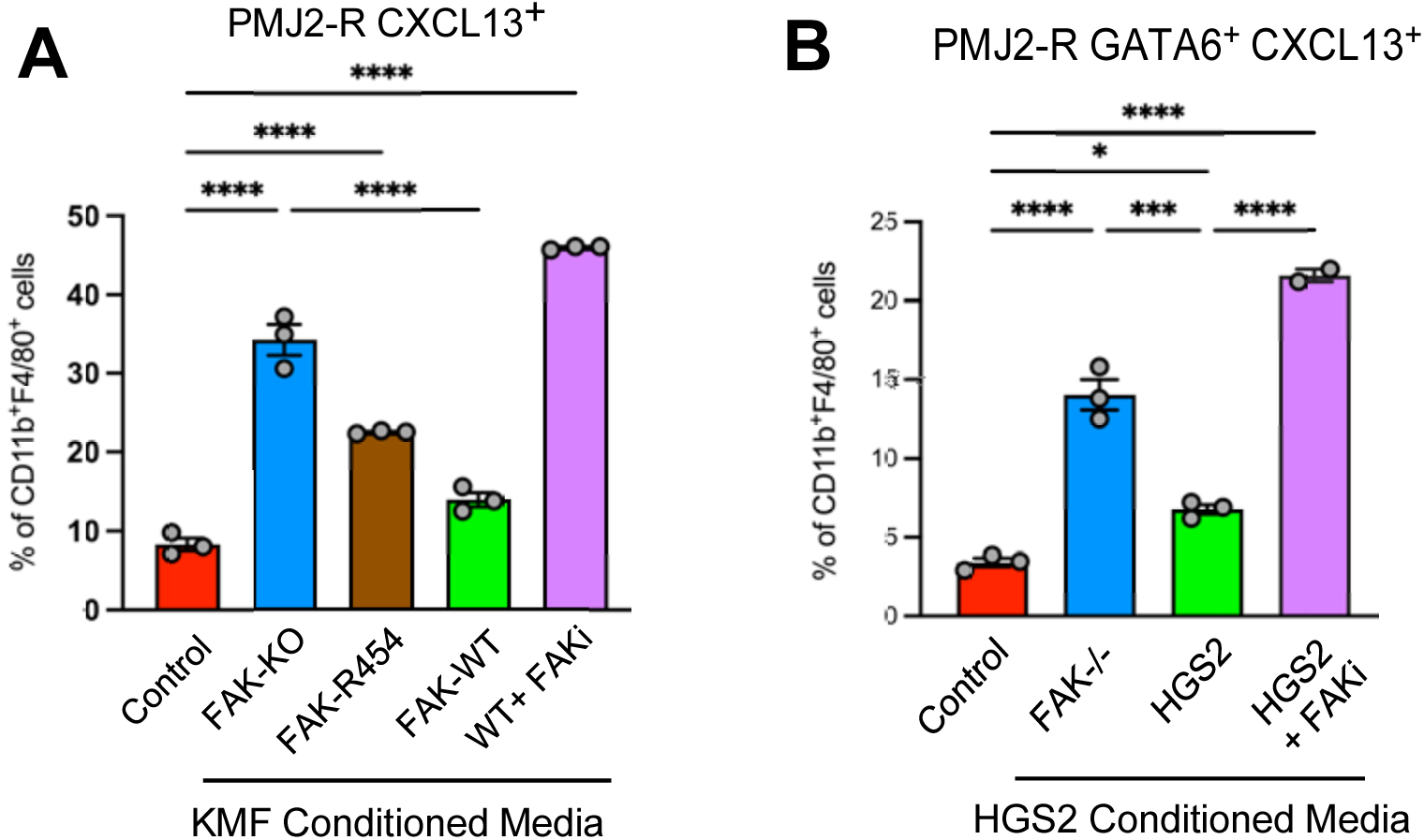
Heat-treated Tumor Conditioned Media (CM) Increases PMJ2-R Macrophage CXCL13. Related to Figure 5. (A) Quantification of CD11b^+^ F4/80^+^ CXCL13^+^ PMJ2-R macrophages by flow cyotometry with the indicated KMF CM incubations. (B) Quantification of CD11b^+^ F4/80^+^ GATA6^+^ CXCL13^+^ PMJ2-R macrophages by flow cyotometry with the indicated HGS2 CM incubations. (B and C) Values are means +/- SEM (*P<0.05, ***P<0.001, ****P<0.0001; one way ANOVA with Tukey’s multiple comparisons test, n=3).

**Figure S5.**
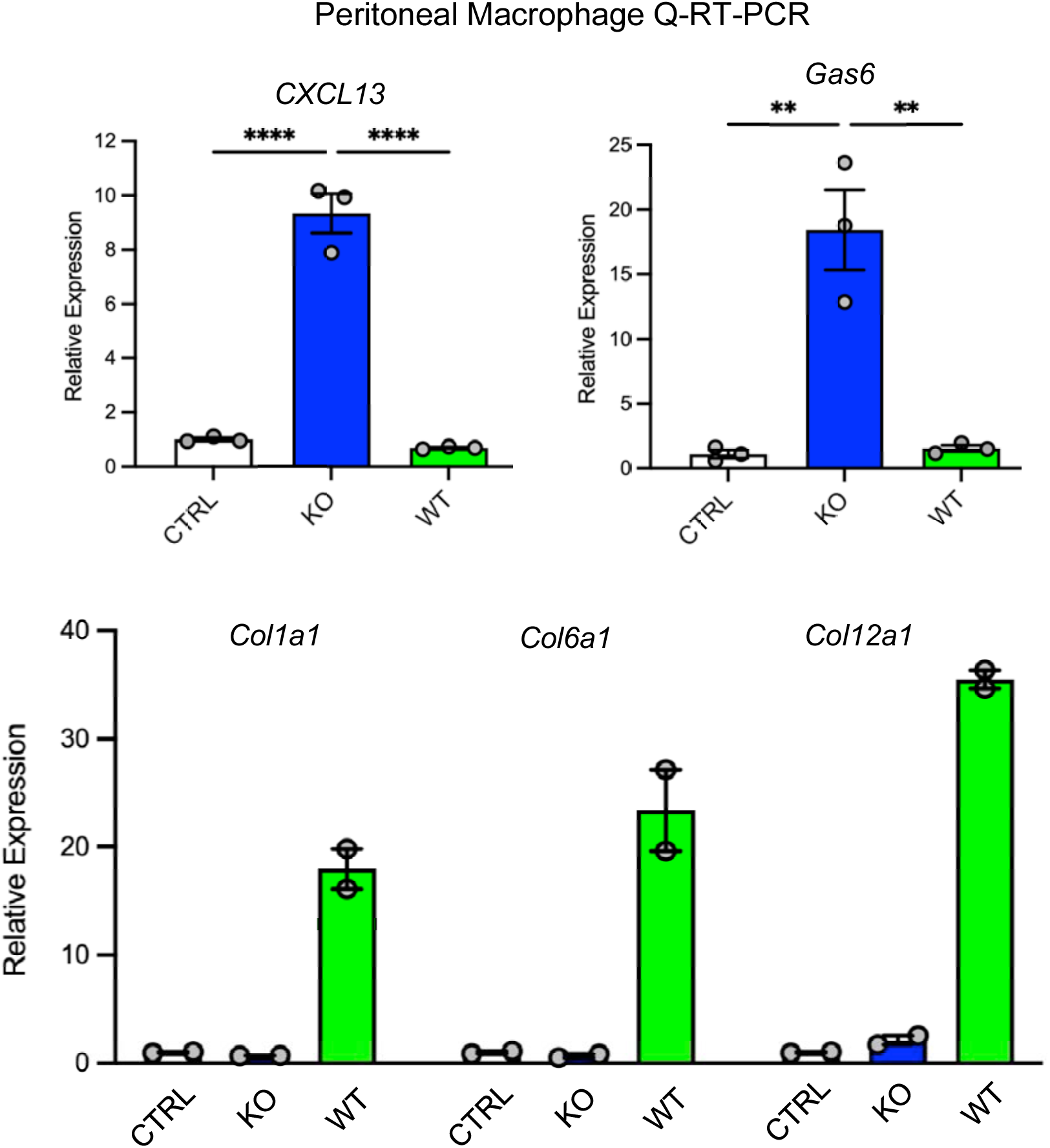
Quantitative Real-time PCR (Q-RT-PCR) of Peritoneal Macrophage mRNA After FAK-KO or FAK-WT Conditioned Media Stimulation. Related to Figure 6. Levels of *Cxcl13, Gas6, Col1a1, Col6a1, and Col12a1* mRNA changes after control (CTRL), KMF FAK-KO, or KMF FAK-WT heat-treated conditioned media stimulation. Values are means +/- SEM, n=2 or 3 independent analyses. (** P<0.01, ****P<0.0001, ANOVA with Tukey’s post-hoc test).

**Figure S6.**
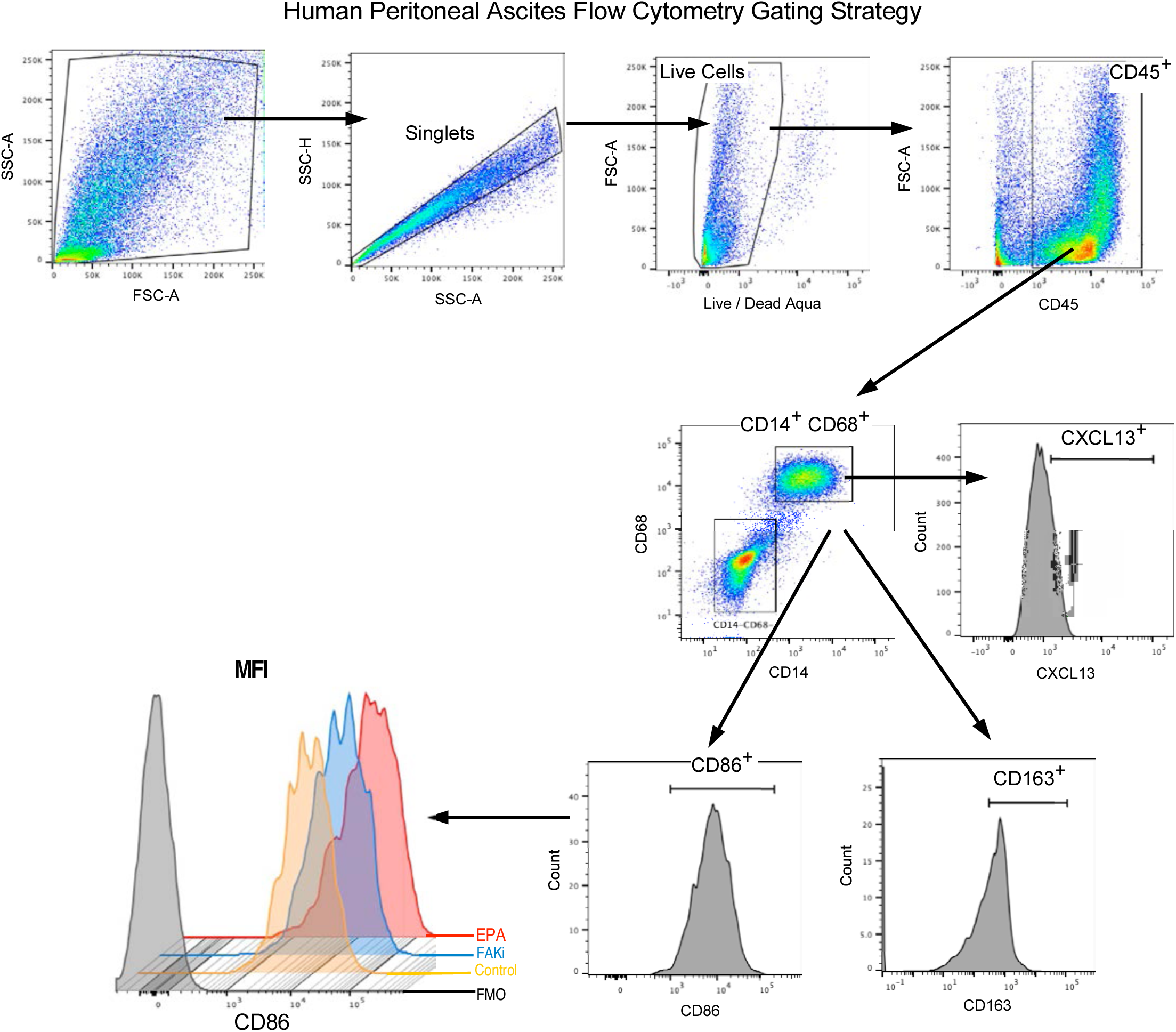
Human Tumor Ascites Flow Cytometry Gating Strategy. Related to STAR Methods Human ovarian tumor and immune cells were collected by paracentesis and CD45^+^, CD14^+^ and CD68^+^ macrophages were analyzed by for changes in CD86^+^ or CD163^+^ levels by counts and mean fluorescence intensity (MFI) upon FAKi or EPA treatment in vitro.

**Table S1.** Single cell RNA sequencing population groupings. Related to Figure 3 and Figure 5.

**Table S2.** HGSOC mRNAs associated with CXCL13. Related to Figure 4.

**Table S3.** Macrophage differential gene expression. Related to Figure 5.

**Table S4.** Lipidomic analysis of whole cell extracts. Related to Figure 7

**Table S5.** Lipidomic analysis of tumor cell conditioned media. Related to Figure 7.

**Table S6.** Lipidomic analysis of purified tumor exosomes. Related to Figure 7.

**Table S7.** Patient Ascites information. Related to Figure 8.

**Table S8.** DNA primers used for PCR and CRISPR. Related to STAR Methods.

## STAR Methods

### Mice

All experimental mouse procedures were reviewed and approved by The University of California San Diego Institutional Animal Care and Use Committee (Protocol S07331). Female C57BL/6 mice (8–10 weeks old) were purchased from Charles River Laboratories. LysMcre GATA6^fl/fl^ YFP^+^ mice (GATA6 WT or KO) on a C57Bl6 background were kindly provided by Gwendalyn Randolph (Washington University). FAK^fl/fl^ mice were used as described^52^. Transgenic mice expressing the transforming region of SV40 T antigen (TAg) under control of the Mullerian inhibitory substance type II receptor (MISIIR) gene promoter were from Denise Connolly (Fox Chase Cancer Center). MISIIR-TAg^+^ and MISIIR-TAg^Low^ mice were developed as described^33,53^. Female NSG (NOD scid gamma) mice (NOD.Cg-*Prkdc^scid^ Il2rg^tm1Wjl^*/SzJ) were purchased from Jackson Labs. C57Bl6 syngeneic HGS2 fallopian tube p53^-/-^ PTEN^-/-^ BRCA2^-/-^ tumor cells (syngeneic with C57Bl6 mice) were from Francis Balkwill (Queen Mary University of London)^54^. Primers used for mouse genotyping at listed (Table S8).

### Cells and Culture

Murine KMF ovarian tumor cells were isolated and cultured as previously described^26^. KMF (*<u>K</u>Ras*, *<u>M</u>yc*, *<u>F</u>AK* amplified) have been exon sequenced and contain DNA gains and losses in gene regions comparable to human HGSOC^26^. KMF FAK knockout (FAK-KO) was created by CRISPR, isolated as a clone (KT13), and exome sequenced as described^26^. KT13 FAK-KO cells were stably reconstituted with wildtype (FAK-WT) or catalytically inactive (FAK-R454) FAK as GFP fusion proteins^26^. KMF cells were cultured in DMEM medium with 10% Fetal Bovine Serum (FBS) and 1% penicillin/streptomycin. HGS2 cells were grown in Advanced DMEM/F-12 medium supplemented with 4% FBS, 1X insulin-transferrin-selenium, 100 ng/ml hydrocortisone, 20 ng/ml EGF, 1X penicillin-streptomycin and L-glutamine. THP-1 cells were cultured in RPMI 1640 medium plus 10% FBS and 1% penicillin/streptomycin. PMJ2-R cells were cultured in modified DMEM medium (ATCC, 30-2002) containing 4 mM L-glutamine, 4500 mg/L glucose, 1 mM sodium pyruvate, 1500 mg/L sodium bicarbonate and 10% FBS. All cell lines were maintained in a humidified incubator at 37°C with 5% CO₂. Mouse ovarian carcinoma (MOVCAR) cells were isolated from ascites of tumor-bearing MISIIR-TAg^+^; FAK^fl/fl^ mice and expanded by growth in non-adherent conditions following methods as described^55^. For 2D growth, MOVCARs were seeded (3 × 10^5^ cells per well) in tissue culture-treated six-well plates (Costar). After 5 days, cells were phase-contrast imaged (Olympus CKX41), collected by trypsinization, and enumerated (ViCell XR, Beckman).

HGS2 FAK and Pyk2 knockout cell lines were generated using CRISPR/Cas9-mediated gene editing using plasmid designed and created by VectorBuilder Inc. Lentiviral plasmids to target murine FAK (*Ptk2*) were pRP[2CRISPR]-Puro-Cas9_D10A-U6>mPTK2[gRNA3] and pRP[2CRISPR]-Puro-Cas9_D10A-U6>mPTK2[gRNA4] with gRNA protospacer sequences GACTCACCTGGGTACTGGCA and ATACTCGTTCCATTGCACCA, respectively. Lentiviral plasmids to target murine PYK2 (*Ptk2b*) pRP[2CRISPR]-Puro-Cas9_D10A-U6>mPTK2b[gRNA1] and pRP[2CRISPR]-Puro-Cas9_D10A-U6>mPTK2b[gRNA2 ] with Grna protospacer sequences GCCCATAGCATTCAGCCAGC and GCTGCACCCACAGATGACCG, respectively. Transfected cells were selected by puromycin treatment (1.5 µg/ml, 36 hours), and single cell clones were isolated by flow cytometry sorting (FACSAria, BD Biosciences). The loss of FAK and Pyk2 protein expression was verified by immunoblotting.

### Immunoblotting

Cells in culture or peritoneal cells from tumor-bearing mice were collected by lavage, washed with cold PBS, whole cell protein lysates were made by RIPA Lysis Buffer (Pierce) addition, and lysates were clarified by centrifugation (16,000 x g, 5 min). Complete™ Mini ETDA-free Protease inhibitor cocktail (Millipore Sigma) and PhoSTOP™ phosphatase inhibitor cocktail (Millipore Sigma) were added Lysis Buffer prior to use. Total protein levels were determined using a bicinchoninic acid assay (Pierce), 25 µg of protein were resolved on Mini-Protean TGX precast gels (4-15% Tris/Glycine gel, BioRad) and transferred to polyvinylidene difluoride membranes using a TransBlot Turbo (Bio-Rad). Immunoreactive protein bands were detected using HRP-conjugated anti-mouse or anti-rabbit antibodies with Clarity Western ECL (enhanced chemiluminescence) substrate reagent and visualized using a ChemiDoc™ Imaging System (Bio-Rad).

### Tumor Growth and Mouse Survival

All animal experiments were performed in accordance with The Association for Assessment and Accreditation for Laboratory Animal Care guidelines and approved by the UCSD Institutional Animal Care and Use Committee (S07331). For tumor implantation studies, cells were transduced with pUltra-Chili-Luciferase (Addgene plasmid #48688) enabling bicistronic expression of dTomato and luciferase and sorted by flow cytometry for equivalent dTomato and GFP expression prior to mouse implantation. For the KMF experiments, unless otherwise indicated, 5 × 10L pUltra-Chili-luciferase-labeled tumor cells were suspended in a 1:1 mixture of culture medium (without FBS) and Matrigel (Corning, 354262), and 200 µL were administered intraperitoneal (IP) injection into 8- to 10-week-old female C57BL/6 mice (Charles River). Mice were randomized into experimental groups on day 5 after tumor cell injection and treated daily with ifebemtinib FAK inhibitor (IN10018, 25 mg/kg, InxMed) by oral gavage. Control group received saline injections. Additional groups were treated with chemotherapeutics including cisplatin (2 mg/kg) plus paclitaxel (10 mg/kg) (CPT), pegylated liposomal doxorubicin (Doxil; 0.15 mg/kg) weekly, or in combinations with the FAK inhibitor. Low endotoxin Anti-TIGIT monoclonal (1B4-mAb; 200 µg) or isotype control (Invivomab, mouse IgG1; Bio X Cell, BE0083) antibody were administered via IP injection at specified time points. Tumor growth was monitored by bioluminescent imaging (IVIS, PerkinElmer).

Tumor-bearing mice were sacrificed at indicated endpoints. Peritoneal tumor and immune cells were harvested by peritoneal lavage using 5 ml of ice-cold buffer (PBS supplemented with 2 mM EDTA and 2% BSA). Cells were processed with red blood cell lysis buffer (BioLegend, 420302), filtered through a 70 µm cell strainer, enumerated, assessed for viability (ViCell XR, Beckman Coulter), and analyzed by flow cytometry and single-cell RNA sequencing. Murine omental and pancreatic tissues were harvested, processed through alcohol and formalin, and paraffin-embedded for histochemical staining. For survival analysis, mice were monitored daily after therapy cessation on day 24 for ascites tumor burden (appearance and body mass increase) and interference of normal behavior (feeding, grooming, lethargy) prior humane euthanasia as defined IACUC guidelines.

MOVCAR orthotopic injection into the ovary-bursa space was performed as described^33^. Briefly, MOVCAR or HGS2 cells were mixed with growth factor reduced Matrigel (Corning), at a concentration of 0.5 × 10^6^ cells per 7 μl and injected into the surgically identified right ovary using a Hamilton syringe, 29.5-gauge needle, and dissecting microscope. Incisions were closed with surgical stables and mice were evaluated for health daily. High resolution ultrasound imaging (Vevo 2100, Visual Sonics) was used to monitor MOVCAR tumor growth in syngeneic TAg^+^ or NSG mice. For MOVCAR tumor metastasis, tumor-bearing NSG mice at 8 weeks were euthanized, primary tumors removed and weighed, lungs were inflated by intratracheal injection, fixed in Bouin’s solution (Sigma), paraffin embedded, sectioned, and stained with hematoxylin and eosin (H&E) for histological evaluation. Tumor metastases per lung lobe was determined by enumerating five sequential sections (evaluated every 25 µm) lung lesions by H&E sections. Mice with >2 MOVCAR lung metastasis were deemed positive.

### Flow Cytometry

Cells were resuspended in flow staining buffer (PBS supplemented with 2 mM EDTA and 2% BSA) and incubated with anti-mouse CD16/32 (BD Biosciences, 553142) or Human TruStain FcX (BioLegend, 422302) Fc-blocking reagent for 15 minutes. For murine samples, surface staining was performed at 4°C for 30 minutes using the following antibodies: LIVE/DEAD Fixable Aqua Dead Cell Stain (Thermo Fisher Scientific, L34957), CD45 (BD Biosciences, 560510), B220 (BioLegend, 103255), TCRβ (eBioscience, 47-5961-82), CD4 (BioLegend, 100509), CD8 (BD Biosciences, 552877), MHC Class II (I-A/I-E; Invitrogen, 50-112-8850), CD11b (BioLegend, 101233), F4/80 (eBioscience, 45-4801-82), CD11c (BioLegend, 117334), CD206 (BioLegend, 141732), and CD155 (BioLegend, 131510). For intracellular staining, cells were fixed and permeabilized using the eBioscience™ Foxp3 / Transcription Factor Staining Buffer Set (eBioscience, 00-5523-00), followed by staining with CXCL13 (eBioscience, 17-7981-82) and GATA-6 (Cell Signaling, 26452). Human peritoneal ascites samples were stained with antibodies against CD45 (BioLegend, 304041) and CD14 (BioLegend, 301803), followed by intracellular staining with CD68 (BioLegend, 333827) and CXCL13/BLC/BCA-1 (R&D Systems, IC8012A-100). Flow cytometry was performed using BD LSR Fortessa and Fortessa X-20 cytometers. Data were analyzed using FlowJo software (version 10).

### Serum Cytokine Quantification

Tumor-bearing mice were exsanguinated by trans cardiac puncture, blood was allowed to clot at for 30 min at RT and centrifuged at 1,000 × g for 10 min to isolate serum. Cytokines and Chemokines were quantified using the LEGENDplex™ Custom Mouse Panel 750 (BioLegend) according to the manufacturer’s instructions.

### Multiplex Immunofluorescence Staining

Murine omental and pancreatic tissues from tumor-bearing mice were harvested, fixed in 10% neutral-buffered formalin, and paraffin-embedded. Tissue sections (5 µm) were baked at 60°C for 1 hour, followed by deparaffinization in xylene (3 × 5 min) and rehydration through successive alcohols (2 × 100%, 2 × 95%, 2 × 70%) into distilled water. Antigen retrieval was performed using citrate-based Antigen Unmasking Solution (Vector Laboratories, H-3300) at 95°C for 30 minutes. Staining was performed using the Intellipath Automated IHC Stainer (Biocare Medical). Endogenous peroxidase activity was quenched using BLOXALL Blocking Solution (Vector Laboratories, SP-6000) for 10 minutes, followed by two washes in TBST (Tris-buffered saline with 0.1% Tween-20). Non-specific binding was blocked with Blocker™ BLOTTO blocking buffer (Thermo, 37530) for 10 minutes.

For multiplex staining with anti-CD3 and anti-B220, sections were incubated with anti-CD3 primary antibody (Abcam, ab16669; 1:500 dilution) for 1 hour, followed by two TBST washes. Detection was performed using HRP-conjugated anti-rabbit polymer (Cell IDx, 2RH-50) for 30 minutes, and developed with Opal 570 fluorophore (Akoya Biosciences, FP1488001KT) for 10 minutes. Slides were washed twice in TBST before proceeding to the second round of antigen retrieval. The second antigen retrieval was performed using the same citrate-based buffer (Vector, H-3300) at 95°C for 30 minutes. After a 5-minute Bloxall block and two TBST washes, sections were incubated with anti-B220 antibody (BD Biosciences, 553086; 1:200 dilution) for 1 hour, followed by two TBST washes. Detection was achieved using HRP-conjugated anti-rat polymer (Cell IDx, 2AH-50) for 30 minutes and developed with Opal 690 fluorophore (Akoya Biosciences, FP1497001KT) for 10 minutes.

For multiplex staining with anti-B220, anti-CD3, anti-F4/80 and anti-CXCL13, sequential rounds of antigen retrieval and antibody staining were performed as described above. The following primary antibodies and detection reagents were used: anti-B220 (BD Biosciences, 553086, 1:200), anti-rat HRP Polymer (Cell IDX, 2AH-50) and Opal 520 (Akoya, FP1487001KT); anti-CD3 (Abcam, ab16669, 1:500), Anti-rabbit HRP Polymer (Cell IDX, 2RH-50), Opal 570 (Akoya, FP1488001KT); anti-F4/80 (Bio-Rad, MCA497BB, 1:50), anti-rat HRP Polymer (Cell IDX, 2AH-50), Opal 620 (Akoya, FP1495001KT); anti-CXCL13 (R&D, AF470, 1:400), anti-goat HRP Polymer (Cell IDX, 2GH-50), Opal 690 (Akoya, FP1497001KT). Each antibody was incubated for 1 hour. Between each staining cycle, slides were washed twice in TBST. All antigen retrieval steps were performed in citrate-based buffer (Vector, H-3300) at 95°C for 30 minutes. Each cycle included peroxidase blocking (5 minutes) prior to primary antibody incubation. After the final staining cycle, sections were washed twice in TBST and twice in distilled water, followed by nuclear counterstaining with DAPI (1μg/mL, 15 minutes). Slides were mounted with VECTASHIELD Vibrance Antifade Mounting Medium (Vector Laboratories, H-1700-10) and cover slipped. Whole-slide images were acquired using the PhenoImager™ Fusion imaging system (Akoya Biosciences).

For MOVCAR tumor staining, FITC-conjugated anti-CD45 antibodies (Invitrogen) at 1 mg/ml in 5% BSA and PBS were incubated for 2 h with FITC-conjugated IgG2b isotype antibodies (Invitrogen) as a negative control. Cell nuclei were visualized by incubation with Heochst 33342 (Invitrogen). Images were sequentially captured at an inverted microscope (IX81; Olympus) at 40X using Hamamatsu ORCA-AG camera, pseudo-colored, overlaid, and merged using Photoshop software. Percent CD45+ staining expressed as percent of nuclei using Image J. For MOVCAR analysis of spontaneous lung metastasis,

### Omental Immune Cell Detection and Quantitation

Omental tissue from ovarian tumor-bearing mice were analyzed by H&E staining and multiplex immunofluorescence on sequential 5 µm tissue sections. TLSs, as represented by clustered co-staining were characterized by the presence of T cells (CD3⁺) and B cells (B220⁺), and their numbers were quantified within defined omental areas. Quantitative image analysis was performed using QuPath (version 0.5.1)^56^. Regions of interest (ROI) (minimum of 3 and maximum of 6 ROIs per tissue section) were manually annotated using the software’s annotation tool. In H&E stained sections, individual cells were identified by optical density SUM, tumor cells and tumor infiltrated immune cells were distinguished by nuclear area and hematoxylin optical density. For multiplex-stained slides, antibodies and nuclear stain were pseudo colored, and color intensities were equalized between different slides by thresholding. Cells were segmented within ROIs based on their nuclear DAPI expression. Single marker measurement classifier was created for each antibody and quantification of positive detection in each ROI was exported. Data is shown as percentage of positive detections to all cells in each ROI, mean of the group +-SEM from at least two individual mice as representative of experimental group.

### Murine Peritoneal Macrophage Isolation

Female C57BL/6 mice (8–12 weeks old) or LysMcre GATA6^fl/fl^ YFP^+^ mice (GATA6 WT or KO) were euthanized in accordance with Institutional Animal Care and Use (IACUC) guidelines. Macrophages were isolated by injecting 5 mL of cold sterile PBS into the peritoneal cavity, the abdomen was gently massaged for 30 seconds to dislodge resident cells, and peritoneal fluid was withdrawn and transferred into sterile 50-mL tubes on ice. Peritoneal fluids from 3–5 mice were pooled to ensure sufficient cell yield. Typically, 2–3 × 10L peritoneal cells per mouse was recovered. Cells were pelleted by centrifugation at 400 × g for 10 minutes at 4°C, the supernatant was discarded, and the cell pellet was resuspended in RPMI-1640 medium supplemented with 10% FBS and 1% penicillin-streptomycin. Cells were seeded onto tissue culture plates and incubated at 37°C (5% CO₂) for 2 hours to allow macrophage adherence. Non-adherent cells were removed by washing 3x with warm PBS and the remaining adherent cells were >90% macrophages. These murine peritoneal macrophages cells were subsequently used for downstream applications.

### Conditioned Medium Collection

Ovarian cancer cell lines (KMF or HGS2) were cultured in their respective complete growth media under standard conditions (37°C, 5% CO₂). Upon reaching ∼70% confluency, cells were washed once with PBS, and the medium was replaced with serum-free medium. After 48 hours, the conditioned medium (CM) was collected, centrifuged at 300 × g for 5 minutes to remove cell debris, and filtered through a 0.22-μm filter (Millipore, SE1M179M6) to ensure sterility. The filtered CM was heat-inactivated at 56°C for 30 minutes in 1 ml aliquots, cooled on ice, and stored at −20°C until further use.

### Transwell Assay

Primary peritoneal macrophages were isolated and seeded into the upper chamber of transwell inserts (8-μm pore size) with KMF FAK-WT or FAK-KO tumor cells in the lower chamber. After 72 h of co-culture, macrophages were harvested and analyzed by flow cytometry. Macrophage population was identified as CD45⁺CD11b⁺F4/80⁺ and assessed for expression of GATA6, CXCL13, or double-positive expression by flow cytometry.

### Peritoneal Macrophage RNA Sequencing

Peritoneal macrophages (PMs) were isolated and cultured in growth medium (RPMI-1640 supplemented with 10% fetal bovine serum and 1% penicillin-streptomycin). After 24 h, 1:1 volume of tumor cell CM and fresh PM growth medium were added. After 72 h, PMs were harvested, and total RNA was extracted using the PureLink RNA Purification Kit (Invitrogen, 12183018A) following the manufacturer’s protocol. RNA library preparation and sequencing was performed by Novogene Inc. (Beijing, China). RNA library preparation was performed using Abclonal mRNA-seq Lib Prep Kit for Illumina® (RK20302) and samples were paired-end 150-bp sequenced using NovaSeq X Plus Series (PE150) platform generating >50 million reads per sample. Reads were aligned to the reference genome using HISAT2. Novogene data pipeline performed differential expression analysis of two conditions/groups by using the DESeq2-R and ClusterProfiler as part of Bioconductor (3.2.1) software. Clustering and grouping analyses used transcripts with FPKM values > 1. Differentially expressed genes were defined as genes with a log_2_ (Fold Change) cutoff.

### Single-cell RNA sequencing

Tumor and immune cells were isolated by peritoneal lavage at experimental day 24 and subjected to single cell 3L RNA sequencing using the 10x Genomics Chromium Next GEM Single Cell 3L HT v3.1 platform. Cell barcoding, reverse transcription, cDNA amplification, and library construction were performed according to the manufacturer’s protocol. Sequencing was carried out on an Illumina NovaSeq 6000 platform, generating approximately 1 billion reads per sample. Demultiplexed FASTQ files were processed using Cell Ranger (v6.1.2, 10x Genomics) for unique molecular identifier (UMI) quantification and alignment to the mm10 mouse reference genome.

Downstream analysis was conducted in R (v4.1.2) using Seurat (v4.3.0) software. Cells were filtered based on the following quality control criteria: 250–10,000 detected genes per cell, 1,000–100,000 UMIs, <20% mitochondrial gene content, and <40% ribosomal gene content. Putative doublets were removed using DoubletFinder (v2.0.3). Data normalization and variance stabilization were performed using SCTransform with regularized negative binomial regression to enhance biological signal. Dimensionality reduction was performed using principal component analysis (PCA), and an integrated shared nearest neighbor (SNN) graph was constructed for clustering. Clusters were identified using the modularity optimization-based SNN clustering algorithm (40 principal components, resolution = 0.4). Differential gene expression analysis across clusters was conducted using the FindAllMarkers function in Seurat, applying the default Wilcoxon rank-sum test. Initial cell type annotation was performed using the ScType method, followed by manual curation based on canonical marker expression.

Raw fastq files were aligned to the prebuilt mouse reference genome (refdata-gex-mm10-2020-A) and quantified using CellRanger v7.1.0. Seurat v5.3.0 was used for downstream analysis. All samples were quality filtered to remove cells with less that 250 features and greater than 10,000 features, overall counts < 40,000, percent mitochondrial reads < 20% and percent ribosomal reads < 40%. Data were transformed using SCTransform with percent mitochondrial reads and percent ribosomal reads as covariates. Samples were integrated and clusters were identified with UMAP visualization at a resolution of 0.8. Doublets were filtered out using DoubletFinder. Cell types were defined by marker genes and informed by predictions from SingleR using the mouse immune cell reference (ImmGenData). Genes of interest were overlaid on the UMAP visualizations with FeaturePlot. Violin plots were made for genes of interest with the VlnPlot function. Code Availability: https://github.com/UCSD-Fisch-Lab/KMF_FAK_single_cell

### TCGA Queries

Co-expression was assessed using Pearson correlation with a two-sided t-test for significance, and adjusted q-values were calculated using FDR correction (Benjamini–Hochberg), using the OVCA_TCGA dataset (www.cBioPortal.org). The top 10 genes were listed in with those cells expressing each gene annotated.

### Quantitative RT-PCR

Total RNA was extracted using the PureLink RNA Mini Kit (Invitrogen, 12183018A) according to the manufacturer’s protocol. cDNA was synthesized using the iScript Reverse Transcription Supermix for RT-qPCR (Bio-Rad, 1708841). Quantitative PCR was performed using iTaq Universal SYBR Green Supermix (Bio-Rad, 1725121) with gene-specific primers and cDNA templates on a CFX Opus 96 Real-Time PCR Systems (Bio-Rad). Gene expression levels were normalized to GAPDH as a housekeeping gene control. Primer sequences used are listed (Table S7).

### Lipidomics

150 µL of cold 5:3:2 MeOH:ACN:H2O (v/v/v) solution was added to the dried cell pellet derived from 5×10^5^ FAK-WT and FAK-KO KMF cells cultured in a murine plasma mimetic medium termed VIMPCS with 1X with physiological supplements ^57^. Samples were vortexed for 30 minutes at 4°C, then centrifuged for 10 minutes at 18k RCF. Using 10 ul injection volumes, the supernatants were analyzed by ultra-high-pressure-liquid chromatography coupled to mass spectrometry (UHPLC-MS - Vanquish and Exploris, Thermo). Metabolites were resolved across a 1.7 um, 2.1 x 150 mm Kinetex C18 column using a 5-minute gradient previously described^58^. 500 µL of cold 5:3:2 MeOH:ACN:H2O (v/v/v) solution was used to reconstitute 40 ul dried media extract. Cold MeOH was added in a 1:25 ratio to the stored media. Samples were then vortexed vigorously for 30 minutes at 4°C, then centrifuged for 10 minutes at 18k RCF. Using 10 uL injection volumes, the supernatants were analyzed by ultra-high-pressure-liquid chromatography coupled to mass spectrometry (UHPLC-MS — Vanquish and Exploris, Thermo). Metabolites were resolved across a 1.7 um, 2.1 x 150 mm Kinetex C18 column using a 5-minute gradient previously described^58^. Using 10 µL injection volumes, non-polar lipids were resolved using UHPLC coupled to ddMS2 using a 5-minute gradient method as previously described^59^.

Following data acquisition, .raw files were converted to .mzXML using RawConverter software. Metabolites were then assigned based on intact mass, ^13^C isotope pattern and retention times in conjunction with the KEGG database and an in-house standard library. Peaks were integrated using Maven (Princeton University). Quality control was assessed as using technical replicates run at beginning, end, and middle of each sequence as previously described. Lipidomics data were analyzed using LipidSearch 5.0 (Thermo Scientific), which provides lipid identification by intact mass, isotopic pattern, and fragmentation pattern to determine lipid class and acyl chain composition.

### Exosome Purification and Analysis

KMF FAK-WT or FAK-KO cancer cells were cultured in six 15-cm culture dishes in DMEM medium supplemented with 10% FBS under standard conditions (37°C, 5% CO₂). Upon reaching 70% confluency, cells were washed twice with 10 mL sterile PBS to remove residual FBS-derived vesicles. Cells were then incubated in serum-free conditioned medium for 48 hours. For exosome isolation, conditioned media was transferred to 50-mL conical tubes and centrifuged at 500 × g for 15 minutes at 4°C to remove cells. The resulting supernatants were centrifuged at 10,000 rpm for 20 minutes at 4°C to remove cell debris and large vesicles. Supernatants were transferred into SW28 tubes and exosomes were pelleted by ultracentrifugation at 110,000 × g for 70 minutes at 4°C (Beckman Coulter, SW28 rotor). Supernatants were discarded carefully. Pellets were resuspended in sterile PBS and combined into a single SW28 tube. A second ultracentrifugation was performed at 110,000 × g for 70 minutes at 4°C to wash the vesicles. Following centrifugation, supernatants were discarded, and the final exosome pellets (from 6 plates) were resuspended in 200 µL of PBS. The size distribution and concentration of isolated exosomes were determined using Nanoparticle Tracking Analyzer (NTA) on a ZetaView instrument (Particle Metrix) according to the manufacturer’s instructions.

### Human Patient Ascites

Newly diagnosed or recurrent ovarian cancer patients with ascites were consented and materials from paracentesis were collected by the Department of Pathology, University of California School of Medicine, Moores Cancer Center, USA under Institutional Review Board approved protocol (IRB 181755). All available patient samples were evaluated by experienced pathologists for confirmation of histological type and biomarkers as indicated (Table S6). Peritoneal malignant ascites was centrifuged at 300 × g for 10 minutes at 4°C. Cell pellets were treated with red blood cell lysis buffer (BioLegend, 420302) for 5 minutes at room temperature, followed by the addition of PBS to terminate the reaction. Intact cells were pelleted by centrifugation and resuspended in growth medium containing RPMI 1640 supplemented with 10% heat-inactivated FBS, 2 mM L-glutamine, and 1% penicillin-streptomycin.

Cell suspensions were washed by repeated centrifugation and filtered through a 70 µm cell strainer. Total cell counts were obtained using a ViCell XR automated hemocytometer (Beckman Coulter). Collected cells were either frozen, directly used for experiments, analyzed by flow cytometry, or purified by density gradient separation. Crude patient ascites cell suspensions, which contain both tumor and immune cells, were seeded in 6-well poly-HEMA-coated plates (Corning, 3471). Cells were cultured for 48 hours, and 2 × 10L cells per well were treated with either ifebemtinib FAK inhibitor (IN10018, 1 µM), eicosapentaenoic acid (EPA, 50 µM), or docosahexaenoic acid (DHA, 50 µM) for 48 h, and analyzed by flow cytometry.

### Purification of Human Tumor Associated Macrophages

To isolate macrophages from patient ascites, a discontinuous Percoll gradient was prepared as previously described^60^. Briefly, 10 mL of 70% Percoll, 15 mL of 45% Percoll, 20 mL of 25% Percoll (containing the cell suspension), and 5 mL of PBS were layered sequentially into a 50 mL conical tube and centrifuged at 800 × g for 30 minutes at room temperature without braking. Macrophage-enriched cells were collected from the 25–45% Percoll interface. Fractions were washed with growth medium and centrifuged at 800 × g for 7 minutes. To further enrich for macrophages, cells from the 25–45% Percoll interface were plated in tissue culture dishes and incubated for 2 hours at 37°C. Non-adherent cells were gently removed by washing twice with warm PBS, leaving a population enriched for adherent macrophages.

### Statistics

Unless indicated, results presented are from at least two independent experiments. Statistical analyses were performed in Prism v10 (GraphPad Software). For experimental groups of three or more, statistical significance was calculated based on one-way ANOVA with Tukey’s multiple comparison test. Unpaired T test was used to determine statistical difference between the means from two different samples. P values <0.05 were considered significant.

### Sex as a Biological Variable

These studies focused on ovarian cancer, a disease that biologically affects females only. Accordingly, all in-vitro and in-vivo experiments involved female-derived cell lines or animal models where applicable. The OV-TCGA (Ovarian Serous Cystadenocarcinoma, The Cancer Genome Atlas) patient dataset is de-identified and publicly available.

---

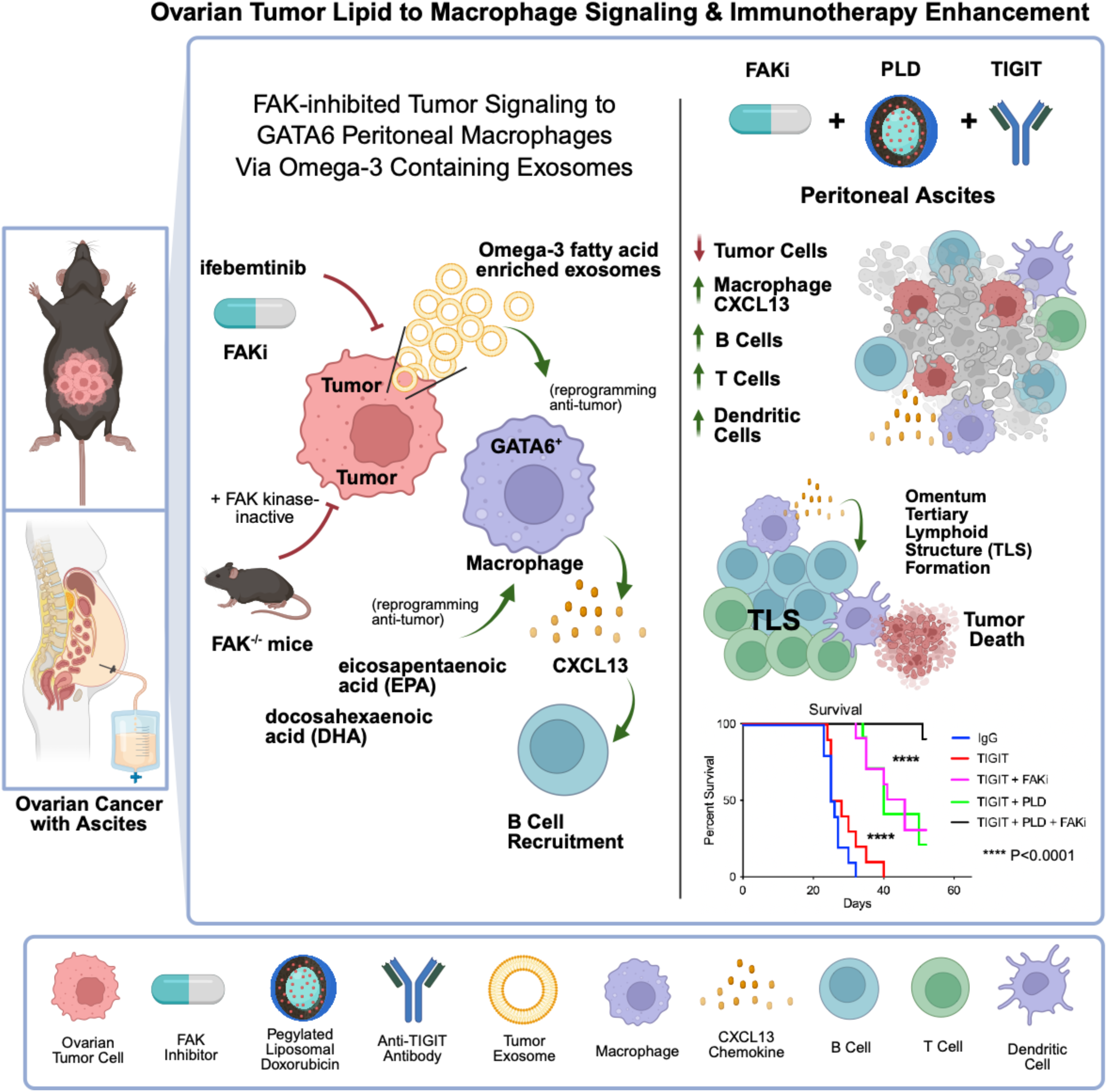

