## Supplementary material for "Ovarian Tumor FAK Inhibition Releases Omega-3 Fatty Acids Stimulating GATA6 Peritoneal Macrophage CXCL13 Production Enhancing Immunotherapy": Key Resource Table

### TABLE FOR AUTHOR TO COMPLETE

***Please do not add custom subheadings.*** *If you wish to make an entry that does not fall into one of the subheadings below, please contact your handling editor or add it under the “other*” subheading*.* ***Any subheadings not relevant to your study can be skipped.*** *(****NOTE:*** *references should be in numbered style, e.g., Smith et al.^1^)*

**Key resources table**

| **REAGENT or RESOURCE** | **SOURCE** | **IDENTIFIER** |
| --- | --- | --- |
| Antibodies | | |
| Alexa Fluor® 700 Rat Anti-Mouse CD45 | BD Biosciences | Cat# 560510 |
| APC anti-mouse CD155 (PVR) Antibody | BioLegend | Cat# 131510 |
| LIVE/DEAD™ Fixable Aqua Dead Cell Stain Kit | Thermo | Cat# L34957 |
| Brilliant Violet 711™ anti-mouse/human CD45R/B220 Antibody | BioLegend | Cat# 103255 |
| TCR beta Monoclonal Antibody (H57-597), APC-eFluor™ 780 | eBioscience | Cat# 47-5961-82 |
| FITC anti-mouse CD4 Antibody | BioLegend | Cat# 100509 |
| BD Pharmingen™ PE-Cy™7 Rat Anti-Mouse CD8a | BD Biosciences | Cat# 552877 |
| Brilliant Violet 605™ anti-mouse CD11c Antibody | BioLegend | Cat# 117334 |
| Brilliant Violet 570™ anti-mouse/human CD11b Antibody | BioLegend | Cat# 101233 |
| F4/80 Monoclonal Antibody (BM8), PerCP-Cyanine5.5 | eBioscience | Cat# 45-4801-82 |
| CXCL13 Monoclonal Antibody (DS8CX13), APC | eBioscience | Cat# 17-7981-82 |
| GATA-6 (D61E4) XP® Rabbit mAb (PE Conjugate) | Cell Signaling | Cat# 26452 |
| MHC Class II (I-A/I-E) Monoclonal Antibody (M5/114.15.2) | eBioscience | Cat# 50-112-8850 |
| PE/Dazzle™ 594 anti-mouse CD206 (MMR) Antibody | BioLegend | Cat# 141732 |
| Brilliant Violet 605™ anti-human CD45 Antibody | BioLegend | Cat# 304041 |
| FITC anti-human CD14 Antibody | BioLegend | Cat# 301803 |
| Brilliant Violet 421™ anti-human CD68 Antibody | BioLegend | Cat# 333827 |
| Human CXCL13/BLC/BCA-1 APC-conjugated Antibody | R&D Systems | Cat# IC8012A-100 |
| Human TruStain FcX™ (Fc Receptor Blocking Solution) | BioLegend | Cat# 422302 |
| BD Pharmingen™ Purified Rat Anti-Mouse CD16/CD32 (Mouse BD Fc Block™) | BD Biosciences | Cat# 553142 |
| Anti-TIGIT [1B4] VivopureX | Absolute Antibody | Cat# Ab01258-1.1-VXB |
| InVivoMAb mouse IgG1 isotype control | Bio X Cell | Cat# BE0083 |
| Anti-FAK, clone 4.47 | Millipore Sigma | Cat# 05–537 |
| Phospho-FAK (Tyr397) Monoclonal Antibody (141-9) | Thermo | Cat# 44-625G |
| Pyk2 (5E2) Mouse mAb | Cell Signaling | Cat# 3480 |
| E-Cadherin (24E10) Rabbit mAb | Cell Signaling | Cat# 3195 |
| N-Cadherin (13A9) Mouse mAb | Cell Signaling | Cat# 14215 |
| Beta Actin Monoclonal antibody | Proteintech | Cat# 60008-1-Ig |
| β-Tubulin (9F3) Rabbit mAb | Cell Signaling | Cat# 2128 |
| Anti-CD3 epsilon antibody [SP7] | Abcam | Cat# ab16669 |
| BD Pharmingen™ Biotin Rat Anti-Mouse CD45R/B220 | BD Biosciences | Cat# 553086 |
| Mouse CXCL13/BLC/BCA-1 Antibody | R&D systems | Cat# AF470 |
| Rat anti Mouse F4/80:Biotin | Bio-Rad | Cat# MCA497BB |
| UltraPolymer Goat anti-Rabbit IgG (H&L) – HRP | Cell IDx | Cat# 2RH-050 |
| UltraPolymer Goat anti-Rat IgG (H&L) – HRP | Cell IDx | Cat# 2AH-050 |
| UltraPolymer Donkey anti-Goat IgG (H&L) – HRP | Cell IDx | Cat# 2GH-050 |
| VIMPCS - murine plasma mimetic medium | Wisentbioproducts | Cat# 319-268-cl |
| IgG2b isotype antibodies | Invitrogen |  |
| Antigen Unmasking Solution | Vector Laboratories | Cat# H-3300 |
| BLOXALL Blocking Solution | Vector Laboratories | Cat# SP-6000 |
| BLOTTO blocking buffer | Thermo | Cat# 37530 |
| HRP-conjugated anti-rabbit polymer | Cell IDx | Cat# 2RH-50 |
| Opal 570 fluorophore | Akoya Biosciences | Cat# FP1488001KT |
| citrate-based buffer | Vector | Cat# H-3300 |
| HRP-conjugated anti-rat polymer | Cell IDx | Cat# 2AH-50 |
| Opal 690 fluorophore | Akoya Biosciences | Cat# FP1497001KT |
| Opal 520 fluorophore | Akoya Biosciences | Cat# FP1487001KT |
| Opal 620 fluorophore | Akoya Biosciences | Cat# FP1495001KT |
| VECTASHIELD | Vector Laboratories | Cat# H-1700-10 |
| Growth factor reduced Matrigel | Corning | Cat# 354262 |
| Bouin’s solution | Sigma | Cat# HT10132 |
| Red blood cell lysis buffer | BioLegend | Cat# 420302 |
| Luciferin |  |  |
| **Bacterial and virus strains** | | |
| Ad-Cre-GFP | Vector Biolabs | Cat# 1700 |
| **Biological samples** |  |  |
| Human ascites from paracentesis | This paper |  |
| **Chemicals, peptides, and recombinant proteins** | | |
| RPMI 1640 | Thermo | Cat# 11875093 |
| DMEM | Thermo | Cat# 11995065 |
| Modified DMEM | ATCC | Cat# 30-2002 |
| Advanced DMEM/F-12 | Thermo | Cat# 12634010 |
| Fetal bovine serum | R&D Systems | Cat# S12450 |
| Insulin/transferrin/selenium | Invitrogen | Cat# 51300 |
| hydrocortisone | Sigma | Cat# H0135 |
| Murine Epidermal Growth Factor (EGF) | Sigma | Cat# E4127 |
| RBC Lysis Buffer (10X) | Biolegend | Cat# 420302 |
| RIPA Lysis and Extraction Buffer | Thermor | Cat# 89900 |
| Mouse IL-4 Recombinant Protein | PeproTech | Cat3 214-14 |
| Lipopolysaccharides (LPS) | Sigma-Aldrich | Cat# L6529 |
| Mouse M-CSF Recombinant Protein | PeproTech | Cat3 315-02 |
| Eicosapentaenoic Acid (EPA) | MCE | Cat# HY-B0660 |
| Linolenic Acid (LA) | Thermo | Cat# 21504 |
| Docosahexaenoic Acid (DHA) | Cayman Chemical | Cat# 90310 |
| PROTAC Degrader FC11 | Tocris Bioscience | Cat# 7306 |
| DOXOrubicin HCI Liposome Injection | Dr.Reddy’s | NDC 43598-0283-35 |
| Paclitaxel | Pfizer | NDC 61703-0342-09 |
| Cisplatin | West ward | NDC 0143-9504-01 |
| Foxp3 / Transcription Factor Staining Buffer Set | eBioscience | Cat# 00-5523-00 |
| Antigen Unmasking Solution, Citrate-Based | Vector Laboratories | Cat# H-3300-250 |
| BLOXALL® Endogenous Blocking Solution | Vector Laboratories | Cat# SP-6000-100 |
| Blocker™ BLOTTO in TBS | ThermoFisher | Cat# 37530 |
| Opal 570 Reagent Pack | Akoya Biosciences | Cat# FP1488001KT |
| Opal 690 Reagent Pack | Akoya Biosciences | Cat# FP1497001KT |
| Opal 520 Reagent Pack | Akoya Biosciences | Cat# FP1487001KT |
| Opal 620 Reagent Pack | Akoya Biosciences | Cat# FP1495001KT |
| VECTASHIELD Vibrance® Antifade Mounting Medium | Vector Laboratories | Cat# H-1700-10 |
| Ifebemtinib (IN10018) FAK Inhibitor | InxMed | clinical compound |
| IVISbrite D-Luciferin Potassium Salt Bioluminescent Substrate | Revvity | Cat# 122799 |
| Corning® Matrigel® Matrix High Concentration (HC), Phenol-Red Free, LDEV-free | Corning | Cat# 354262 |
| Heochst 33342 | Thermo | Cat# 62249 |
| Mini ETDA-free Protease inhibitor cocktail | Sigma | Cat#11836170001 |
| PhoSTOP™ phosphatase inhibitor cocktail | Millipore | Cat# 4906845001 |
| Clarity Western ECL | BioRad | Cat# 1705060S |
| **Critical commercial assays** | | |
| eBioscience™ Foxp3 / Transcription Factor Staining Buffer Set | eBioscience | Cat# 00-5523-00 |
| LEGENDplex Custom Mouse Panel 750 | Biolegend | Cat# 900001850 |
| PureLink RNA Mini Kit | Invitrogen | Cat# 12183018A |
| iScript Reverse Transcription Supermix for RT-qPCR kit | Bio-Rad | Cat# 1708841 |
| iTaq Universal SYBR Green Supermix | Bio-Rad | Cat# 1725121 |
| Abclonal mRNA-seq Lib Prep Kit | Illumina | Cat# RK20302 |
| Transwell chambers (8 µm) | Costar |  |
| 0.22-μm filter | Millipore | Cat# SE1M179M6 |
| Pierce™ BCA Protein Assay Kits | ThermoFischer | Cat# 23225 |
| Mini-PROTEAN® TGX™ Precast Gels | BioRad | Cat# 4561033 |
| Corning™ Costar™poly-HEMA-coated 24-well plates | Fischer Scientific | Cat# 09-761-146 |
| **Deposited data** | | |
| Mouse bulk RNA sequencing of macrophage | This paper | GEO: |
| Mouse ovarian cancer single cell RNA sequencing | This paper | GEO: |
| **Experimental models: Cell lines** | | |
| KMF (ID8-IP) | PMID: 23275034 | Schlaepfer Lab |
| THP-1 | ATCC | Cat# TIB-202 |
| HGS2 | Ximbio | Cat# 160538 |
| PMJ2-R | ATCC | Cat# CRL-2458 |
| 293T | ATCC | Cat# CRL-3216 |
| MOVCAR | This paper |  |
| **Experimental models: Organisms/strains** | | |
| Mouse: C57BL/6 | Charles River Laboratories | C57BL/6 Mouse |
| Mouse: FAK fl/fl | PMID: 15967814 |  |
| Mouse: TAg | PMID: 12649204 |  |
| Mouse: LysMcre GATA6fl/fl YFP+ mice | PMID: 25024137 |  |
| **Oligonucleotides** | | |
| (see table) |  |  |
| **Recombinant DNA** | | |
| pUltra-Chili-luciferase | Addgene | Cat# 48688 |
| **Software and algorithms** | | |
| FlowJo 10 | Becton-Dickinson | https://www.flowjo.com/ |
| Prism 10 | Graphpad | https://www.graphpad.com/ |
| QuPath-0.5.0-x64 | PMID: 29203879 | https://qupath.readthedocs.io/en/latest/index.html |
| NovoMagic | Novogene | https://cssamerica.novogene.com/pub/novoMagic |
| Python | Python Software Foundation | [https://www.python.org](https://www.python.org/) |
| ImageJ | PMID: 22930834 | https://imagej.nih.gov/ij/ |
| RawConverter |  |  |
| Maven |  |  |
| LipidSearch 5.0 | thermofisher.com | Cat# OPTON-30880 |
| R (v4.1.2) |  |  |
| Seurat (v4.3.0) |  |  |
| DoubletFinder (v2.0.3) |  |  |
| SCTransform |  |  |
| CellRanger v7.1.0 | 10xgenomics.com/software |  |
| FeaturePlot |  |  |
| SingleR |  |  |
| Adobe Photoshop 2025 | Adobe.com | https://www.adobe.com/products/photoshop.html |
| Canvas X Draw (v7.1 build 7112) | canvasgfx.com | https://www.canvasgfx.com/products/canvas-x-draw |
| **Other** | | |
| mm10 mouse reference genome |  |  |
| mouse immune cell reference (ImmGenData) |  |  |
